# *Sox17* expression in endocardium precursor cells regulates heart development in mice

**DOI:** 10.1101/548289

**Authors:** Rie Saba, Keiko Kitajima, Lucille Rainbow, Sylvia Engert, Mami Uemura, Hidekazu Ishida, Ioannis Kokkinopoulos, Yasunori Shintani, Shigeru Miyagawa, Yoshiakira Kanai, Masami Azuma-Kanai, Peter Koopman, Chikara Meno, John Kenny, Heiko Lickert, Yumiko Saga, Ken Suzuki, Yoshiki Sawa, Kenta Yashiro

**Author notes:** To whom correspondence may be addressed. Kenta Yashiro, MD, PhD, Department of Anatomy, Division of Anatomy and Developmental Biology, Kyoto Prefectural University of Medicine, 465 Kajii-cho, Kawaramachi-Hirokouji, Kamigyo-ku, Kyoto 602-8566, Japan.

## Abstract

The endocardium is the endothelial component of the vertebrate heart and plays a key role in heart development. Cardiac progenitor cells (CPCs) that express the homeobox gene *Nkx2-5* give rise to the endocardium. Where, when, and how the endocardium segregates during embryogenesis have remained largely unknown, however. We now show that *Nkx2-5*^+^ CPCs that express the Sry-type HMG box gene *Sox17* specifically differentiate into the endocardium in mouse embryos. Approximately 20% to 30% of *Nkx2-5*^+^CPCs transiently express *Sox17* from embryonic day (E) 7.5 to E8.5.Although *Sox17* is not essential or sufficient for endocardium fate, it can bias the fate of CPCs toward the endocardium. On the other hand, *Sox17* expression in the endocardium is required for heart development. Deletion of *Sox17* specifically in the mesoderm markedly impaired endocardium development with regard to cell proliferation and behavior. The proliferation of cardiomyocytes, ventricular trabeculation, and myocardium thickening were also impaired in a non–cell-autonomous manner in the *Sox17* mutant, resulting in anomalous morphology of the heart, likely as a consequence of down-regulation of NOTCH signaling. Changes in gene expression profile in both the endocardium and myocardium preceded the reduction in NOTCH-related gene expression in the mutant embryos, suggesting that *Sox17* expression in the endocardium regulates an unknown signal required for nurturing of the myocardium. Our results thus provide insight into differentiation of the endocardium and its role in heart development.

**Significance:** The endocardium is vital for vertebrate heart development; however, the molecular mechanisms regulating fate determination and differentiation remain largely unknown. Here, we show that a part of the earliest cardiac progenitor cells (CPCs) transiently and exclusively express *Sry*-type HMG box gene *Sox17* in the mouse embryo. *Sox17*-expressing CPCs specifically differentiate to the endocardium. *Sox17* biases the fate of CPCs toward the endocardium, and regulates proliferation and cellular behavior cell autonomously. Conversely, *Sox17* in the endocardium regulates the myocardium non-cell autonomously. Notably, *Sox17* is required for the ventricular trabeculation via the NOTCH signal that is not directly induced but maintained by *Sox17*. This study, thus, sheds light on endocardium development.

## Introduction

Heart development is one of the earliest events of vertebrate organogenesis. Cardiac progenitor cells (CPCs) give rise to the myocardium, endocardium, epicardium, smooth muscle, fibroblasts, and endothelium of coronary vessels in the mammalian heart. In the mouse embryo, CPCs originate between embryonic day (E) 6.25 and E7.5 from nascent mesoderm cells in the primitive streak that express the basic helix-loop-helix (bHLH) transcription factor gene *Mesp1*. These mesoderm cells populate the heart field as CPCs at the most anterior region of the embryo, and they begin to express CPC markers at the early allantoic bud (EB) stage (E7.5). In addition to well-validated CPC marker genes encoding transcription factors—including *Nkx2-5, Isl1*, and *Tbx5* (1–3)—analysis of alternative markers has provided the basis for a model of the cell lineage tree and the mechanics of cardiomyocyte differentiation (4, 5). Among the cell types constituting the heart, the development of endocardial cells has remained largely uncharacterized.

The endocardium is the innermost layer and endothelial component of the heart. During heart development, the endocardium provides a source of cells for the valves, the membranous portion of the ventricular septum, the endothelium of coronary vessels, and cardiac fibroblasts. It plays a key role in ventricular trabeculation, myocardial compaction, as well as valve and coronary vessel formation. In the mouse embryo, endocardial cells are first recognized morphologically as a proendocardium layer between the myocardium and definitive endoderm layers at the one- to two-somite stage, when expression of the transcription factor gene *Nfatc1*, an early and unique marker of the endocardium, is initiated (6–8).

In mice, *Mesp1*-expressing nascent mesoderm cells in the primitive streak have been shown to include two types of cell that give rise to the endocardium: unipotent endocardium precursors, and bipotent cardiac progenitors that contribute to cardiomyocytes and the endocardium (9). No molecular marker has been identified to distinguish these two cell types from each other, but both likely become *Nkx2-5*– positive CPCs (1, 10). Downstream of *Mesp1*, the ETS-related transcription factor gene *Etv2* (also known as *Er71* or *Etsrp71*) plays a role in endocardium development at the top of the endothelial genetic cascade (11). Disruption of *Etv2* was found to result in loss of the endocardium (12, 13), suggesting that the endocardium originates from *Etv2*^+^mesoderm. It remains unclear, however, whether *Etv2* determines endocardium fate or only confers competence for endocardium differentiation. Downstream of *Etv2*, a network of transcription factors—including ETS, SOX, GATA, and RBPJκ—regulates endocardium differentiation. These factors likely activate expression of *Flk1* and *Dll4* in the endocardium (14, 15), with these genes being implicated in vascular endothelial growth factor (VEGF) and NOTCH signaling essential for endocardium development (16, 17). However, the network of transcription factors responsible for the induction of endocardium fate remains largely unidentified.

Among three Sry-type HMG box F (SOXF) transcription factor genes—*Sox7, Sox17*, and *Sox18*—expression of *Sox17* was long regarded as specific for the endoderm including the visceral and definitive endoderm (18). However, *Sox17* was subsequently shown to be essential for vascular development and definitive hematopoiesis (19, 20). Lineage tracing for *Sox17*-expressing cells revealed that they contribute not only to the endoderm but also to the mesoderm that gives rise to the endocardium (21). Conditional knockout of *Sox17* in the endothelial lineage of mice resulted in the loss of vascular rearrangement in the embryo and yolk sac as well as in that of definitive hematopoiesis (19, 21). With regard to cardiac development, differentiation of mouse embryonic stem cells into cardiomyocytes in vitro was suppressed in a non–cell-autonomous manner by *Sox17* ablation, although the identity of *Sox17*-expressing cells that are required for cardiomyogenesis has been unclear (22).

Here we show that *Sox17* is expressed in a subset of CPCs during the early phase of mouse cardiac development. Single-cell gene expression profiling revealed that ∼20% to 30% of *Nkx2-5*^+^CPCs are positive for *Sox17* expression. SOX17^+^mesoderm cells were found in the region corresponding to the heart field in E7.5 mouse embryos. This *Sox17* expression is transient, persisting up to E8.5 at the latest. Tracing of *Sox17*^+^cells showed that the endocardium originates from *Sox17*^+^mesoderm cells. Gain-of-function and mesoderm-specific loss-of-function analyses for *Sox17* in mouse embryos revealed that *Sox17* is required for heart development, although it is not essential or sufficient for endocardial fate. Single-cell gene expression profiling for the mesoderm-specific *Sox17* mutant showed that disruption of *Sox17* resulted in misregulation of the transcriptome in endocardial and myocardial cells in a cell-autonomous and non–cell-autonomous manner, respectively. Our findings thus provide insight into development of the endocardium and its relation to heart morphogenesis.

## Results

### *Sox17* Expression in CPCs

We previously performed single-cell gene expression profiling of mouse embryonic CPCs from the EB to early head fold (EHF) stage (23, 24). We validated cell types based on the expression of marker genes including *Sox2* for the epiblast or neural ectoderm, *Sox17* for the endoderm or arterial endothelial cells (18, 25), *Cryptic* (also known as *Cfc1*) for the lateral plate mesoderm (LPM), and *Nkx2-5* or *Tbx5* for CPCs (1, 3). These data revealed that *Sox17* was expressed in 21.9% of *Cryptic*^+^and *Nkx2-5*^+^and/or *Tbx5*^+^ CPCs at the EB stage (Fig. 1*A*). We also confirmed that ∼20% to 30% of CPCs continued to express *Sox17* up to the early somite stage (Fig. 1*A*), suggesting that *Sox17* plays a role in a subset of CPCs.

**Fig. 1.**
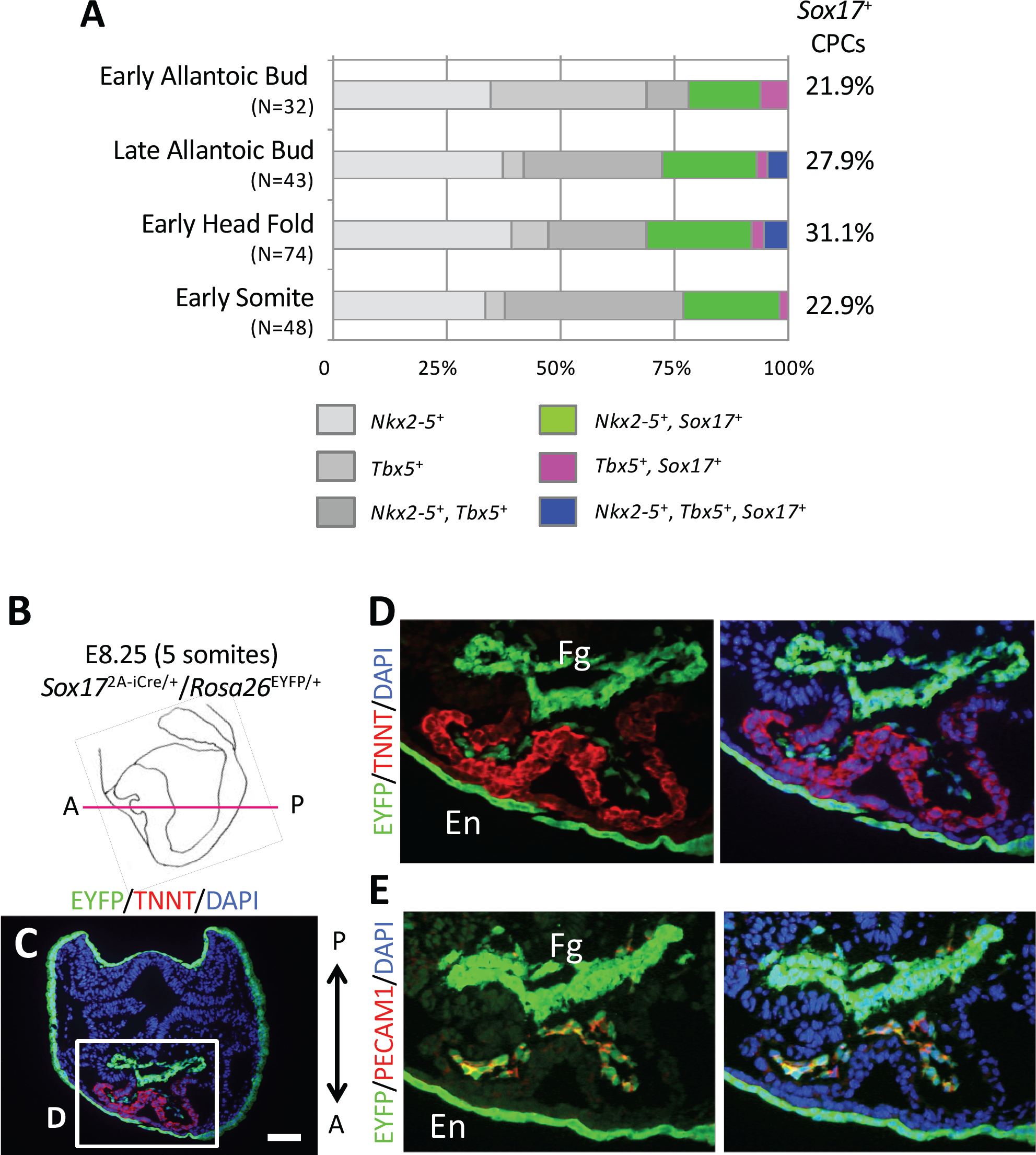
*Sox17*^+^CPCs specific for the endocardium differentiation. (*A*) Proportion of *Sox17*-expressing CPCs in mouse embryos at E7.5 (early allantoic bud, late allantoic bud, and early head fold stages) and E8.5 (early somite stage). *N* values indicate the number of cells examined. The data are derived from our previous study (23). (*B*) Schematic representation of a mouse embryo at the five-somite stage (E8.25) as a left lateral view. The magenta line shows the sectional plane along the anterior (A)– posterior (P) axis in *C*. (*C*–*E*) Immunofluorescence micrographs for EYFP (green), TNNT (red in *C* and *D*), and PECAM1 (red in *E*). Nuclei (blue) were stained with 4′,6-diamidino-2-phenylindole (DAPI). The boxed region in *C* is shown at higher magnification in *D*. The section shown in *E* is adjacent to that in *D*. Fg, foregut; En, endoderm. Scale bar, 100 µm.

We next examined the distribution of SOX17-expressing CPCs in mouse embryos by immunofluorescence analysis (Fig. S1). To distinguish LPM cells from the endoderm, we studied *Mesp1*^Cre/+^/*Rosa26*^EYFP/+^mouse embryos, in which *Mesp1*-expressing mesoderm cells are labeled with enhanced yellow fluorescent protein (EYFP) (26, 27). Whereas SOX17+ cells were rarely detected among the EYFP^+^ LPM cells in embryos at the EB and late allantoic bud stages, SOX17^+^/EYFP^+^ cells were readily apparent at the EHF stage (Fig. S1). At the EHF stage, SOX17^+^ CPCs were marked simultaneously by the CPC marker NKX2-5 in the most anterior portion of the embryo corresponding to the heart field (Fig. S2). As the endoderm layer thickened and the shape of the foregut pocket became more obvious from the late head fold to zero-somite stage (E8.0), SOX17^+^LPM cells became localized more exclusively to a region near the dorsal side of the foregut pocket fold (Fig. S3*A*). From the three-somite stage (E8.25), the number of SOX17^+^ LPM cells decreased concomitantly with the decline in expression of SOX17 in the anterior definitive endoderm (Fig. S3*B*). Only a few SOX17^+^LPM cells were apparent at the sinus venosus. These results thus showed that SOX17 is expressed in CPCs during early embryogenesis.

### Contribution of *Sox17*^+^CPCs to the Endocardium

We next characterized *Nkx2-5*^+^/*Sox17*^+^ CPCs by polymerase chain reaction (PCR) analysis of marker gene expression with single-cell cDNA. Expression of the endothelial marker genes *Dll4* and *Pecam1* was highly correlated with that of *Sox17* in *Nkx2-5*^+^ CPCs at the early somite stage (Fig. S4*A*), whereas the expression of other marker genes, such as *Actn2* (cardiomyocytes) or *Acta2* (smooth muscle cells), was not (Fig. S4*B*) (28–31). These results suggested that *Sox17*-expressing cells contribute to endothelial-like cells, a conclusion also supported by the preferential expression of *Etv2* and *Notch1*, marker genes for arterial endothelial precursor cells, in *Nkx2-5*^+^/*Sox17*^+^CPCs at the early somite stage (>75% for both of *Etv2* and *Notch1*) as well as in *Nkx2-5*^+^CPCs at the EB stage (100% and 72.7% for *Etv2* and *Notch1*, respectively) (Fig. S4*C*) (12, 13, 32). Given that the endothelial component originating from *Nkx2-5*–expressing CPCs becomes the endocardium, it is likely that *Sox17*-expressing CPCs give rise to the endocardium (33).

To examine further whether endocardium cells are indeed derived from *Sox17*-expressing mesoderm cells, we traced the lineage of *Sox17*-expressing cells in *Sox17*^Sox17-2A-iCre/+^/*Rosa26*^EYFP/+^ embryos (Fig. 1*B*–*E*) (21), in which *Sox17*-expressing cells are labeled with EYFP. We found that the progeny of *Sox17*-expressing cells were all PECAM1^+^endocardium cells inside the troponin T (TNNT)^+^myocardium layer at the five-somite stage (E8.5), consistent with the results of a previous study (21). *Sox17*^+^ CPCs are thus the specific precursors of the endocardium.

### *Sox17* Expression Biases CPCs to Express an Endothelial Gene Program

The specificity of *Sox17* expression for the endothelial cell lineage suggested that *Sox17* plays a role in endocardial differentiation of CPCs. We therefore next examined whether *Sox17* expression is sufficient to establish endocardium cell fate in CPCs by forced expression of *Sox17* in *Nkx2-5*–expressing CPCs with the use of a bacterial artificial chromosome (BAC)–based *Nkx2-5*^Sox^17^-IRES-LacZ^ transgene (*Tg*) (Fig. 2*A*). Expression of the *Tg* was confirmed to mimic endogenous *Nkx2-5* expression in CPCs (Fig. 2*B*–*E*). The most severe defect of the *Tg*^+^embryos at E9.5 was anomalous looping and structure of the heart tube, but no obvious morphological abnormalities were apparent before this stage (Fig. 2*F*–*J*). Inside the heart of both severely (Fig. 2*K, L*) and moderately (Fig. S5*A, B*) affected embryos, ventricular trabeculation was impaired. Apoptosis was frequently observed among *Tg*^+^cardiomyocytes (Fig. S5*C*– *E*), whereas the proliferation of Nkx2-5^+^cells inside the heart of *Tg*^+^embryos was not significantly affected (Fig. S5*F*–*H*). The overall gain-of-function phenotype thus suggested that ectopic and excess *Sox17* expression is toxic. Of note, cardiomyocytes expressing PECAM1 were apparent in some *Tg*^+^embryos with a severe phenotype, a phenomenon never observed in wild-type (WT) embryos (Fig. 2*K, L*), suggesting that *Sox17* expression biases the fate of CPCs toward an endothelial-like phenotype but is not sufficient to induce an endocardial cell fate in these cells. This conclusion is consistent with the observation that expression of *Etv2* was not detected in the *Tg*-expressing (*LacZ*-positive) area of *Tg*^+^embryos (Fig. 2*C, E*). On the other hand, *Tg*^+^embryos with a mild to moderate phenotype manifested pronounced aggregation of endocardium cells rather than a monolayer (Fig. S5*B*). The intimate interaction between the endocardium and myocardium was also missing. These findings suggested that *Sox17* expression in the endocardium must be maintained at an appropriate level for proper regulation of cell behavior.

**Fig. 2.**
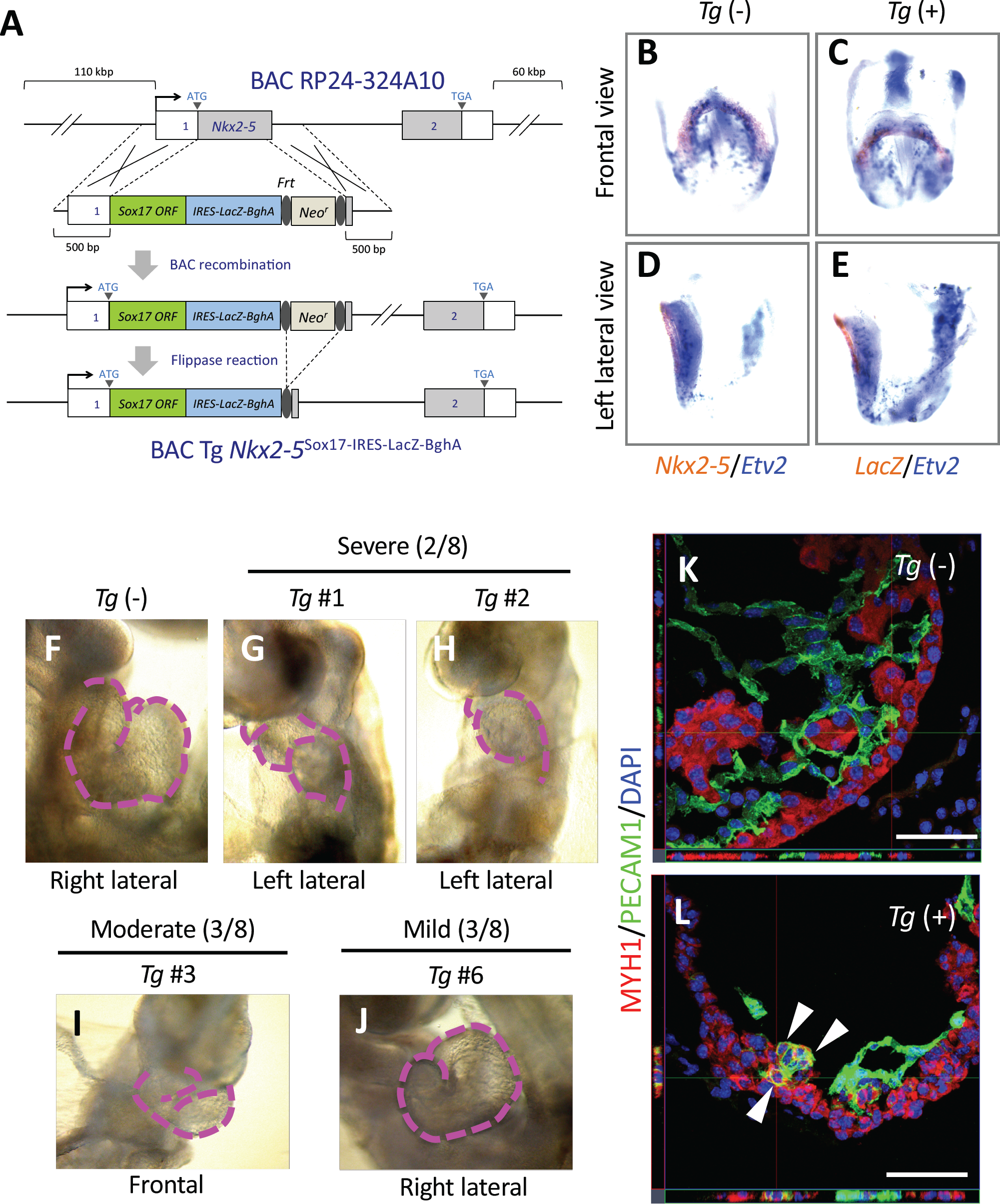
Gain of function of *Sox17* in *Nkx2-5*–expressing CPCs. (*A*) Construction of the BAC-based *Nkx2-5*^Sox17-IRES-LacZ^ *Tg*. (*B*–*E*) Double whole-mount in situ hybridization for *Etv2* (blue) and for either *Nkx2-5* (brown in *B* and *D*) or *LacZ* (brown in *C* and *E*) in E8.25 *Tg*(–) or *Tg*(+) embryos, respectively. (*F*–*J*) The heart of a *Tg*(–) embryo (*F*) and of *Tg*(+) embryos with severe (*G* and *H*), moderate (*I*), or mild (*J*) phenotypes at E9.5. Dashed magenta lines show the outline of the heart tube. (*K* and *L*) Immunofluorescence micrographs for the sarcomere myosin heavy chain MYH1 (red) and PECAM1 (green) in the left ventricle of the heart of E9.5 *Tg*(–) (*K*) and *Tg*(+) (*L*) embryos. Blue, DAPI; Arrowheads, cardiomyocytes expressing PECAM1; Scale bars, 100 µm.

### Cardiac Defects Induced by *Sox17* Deletion in Mesoderm

To elucidate the physiological function of *Sox17* in endocardium development, we deleted *Sox17* specifically in the mesoderm by mating *Mesp1*^Cre/+^mice with mice homozygous for a floxed *Sox17* allele (*Sox17* ^fl/fl^) (19). Although multiple anomalies were apparent in *Mesp1*^Cre/+^/*Sox17*^fl/fl^ embryos at E9.5, no obvious morphological abnormalities were detected before this stage. Growth retardation and ballooning of the pericardial sac were observed after approximately the 20-somite stage, suggestive of a severe defect in the peripheral circulation that likely resulted from embryonic heart failure (Fig. 3*A, B*). Remodeling of the yolk sac vessels was also impaired in the mutant embryos. In WT embryos, the network of capillary vessels that connect to the blood islands, where primitive hematopoiesis gives rise to the production of primary erythrocytes, undergoes remodeling to form the larger vessel network of the yolk sac at E9.5 (Fig. S6). In contrast, such vascular rearrangement did not occur in the mutant. Primitive hematopoiesis appeared almost normal in the mutant at E9.5, consistent with the previous observation that a haematopoiesis defect was only found after E11.5 in *Sox17* mutant (19). These results thus indicated that *Sox17* is essential for vascular development.

**Fig. 3.**
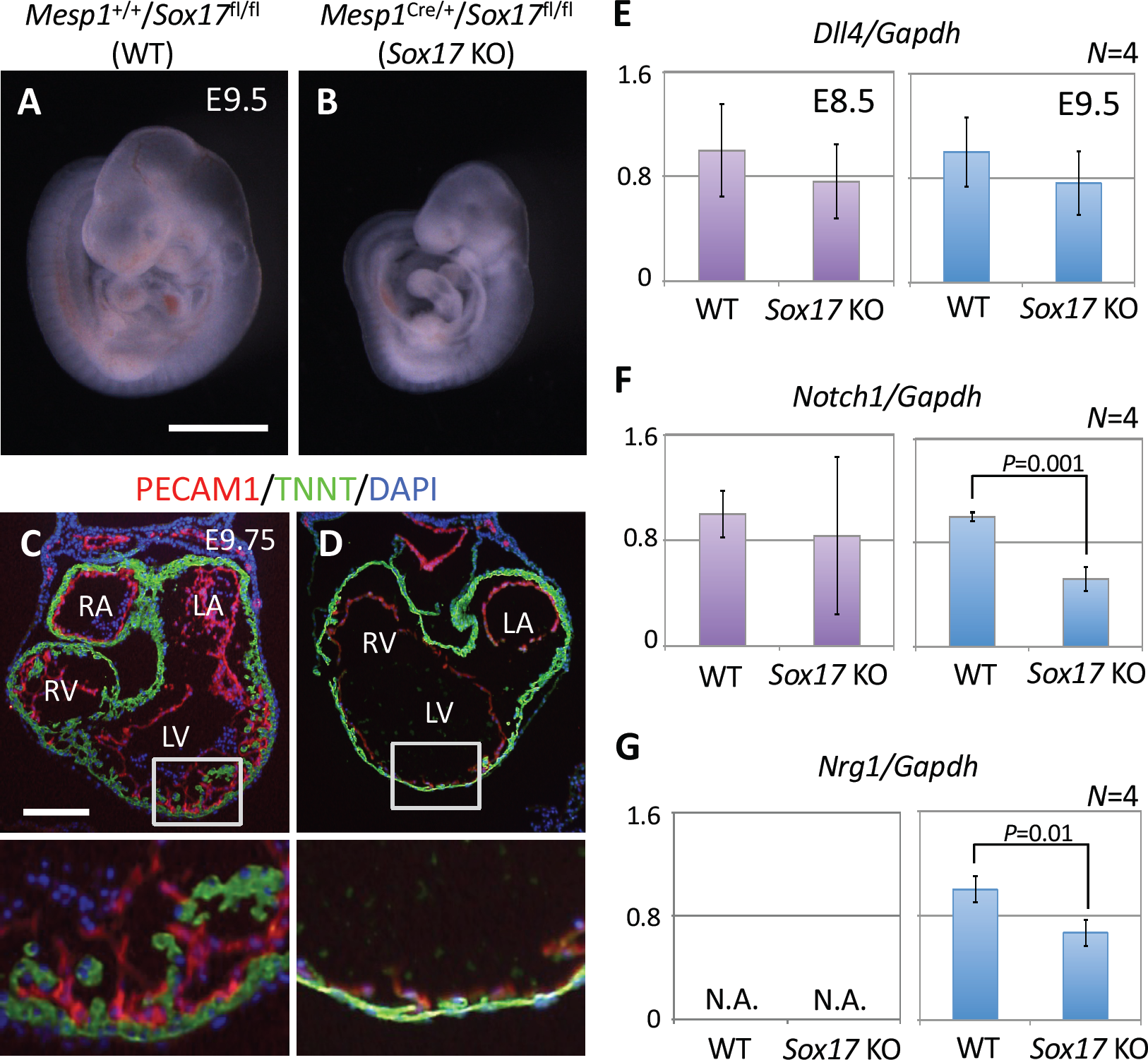
Cardiac defects associated with mesoderm-specific loss of function for *Sox17* in mouse embryos. (*A* and *B*) *Mesp1*^+/+^/*Sox17*^fl/fl^(WT) (*A*) and *Mesp1*^Cre/+^/*Sox17*^fl/fl^(*Sox17* KO) (*B*) embryos at E9.5. Scale bar, 1 mm. (*C* and *D*) Immunofluorescence micrographs in the heart of WT (*C*) and *Sox17* KO (*D*) embryos at E9.75. The boxed regions in the upper panels are shown at higher magnification in the lower panels.Red, PRCAM1; TNNT, Green; Blue, DAPI; LA, left atrium; RA, right atrium; LV, left ventricle; RV, right ventricle. Scale bar, 100 µm. (*E*–*G*) Reverse transcription and real-time PCR analysis of the relative expression levels for the NOTCH signaling– related genes *Dll4* (*E*), *Notch1* (*F*), and *Nrg1* (*G*) in the heart of WT and *Sox17* KO embryos at E8.5 (left panels) and E9.5 (right panels). *Nrg1* expression at E8.5 was below the threshold for detection (N.A., not amplified). Only significant *P* values (Student’s *t* test) are indicated. Means ± SD.

Examination of the heart tube revealed anomalous looping in *Mesp1*^Cre/+^/*Sox17*^fl/fl^embryos at E9.5 (Fig. S7*A*–*F*). In WT embryos, the borders of the outflow tract, ventricle, and atrium were identified by folds with a sharp angle. However, the fold angles were obtuse in the mutant. The endothelial-like phenotype of the endocardium was not impaired in the heart of mutant embryos at E9.75, as revealed by the expression of PECAM1 (Fig. 3*C, D*), indicating that *Sox17* is not essential for endocardium fate in CPCs. However, the number of endocardium cells appeared to be reduced in the mutant. Whereas the endocardium was in intimate contact with the myocardium layer in WT embryos, such “touchdown” sites (34, 35) were far fewer in the mutant. The size of cardiomyocytes was also reduced in the mutant embryos. Importantly, myocardial trabeculation was severely affected and the ventricular wall was markedly thinner in the mutant. Together, these observations indicated that the physiological function of the endocardium was so anomalous in the *Sox17* mutant that maturation of the myocardium was affected in a non–cell-autonomous manner.

We next examined the expression levels of NOTCH signaling–related genes in the developing heart, given that a *Dll4-Notch1-Nrg1* pathway plays an important role in the differentiating endocardium as well as in ventricular trabeculation (36). The expression level of *Dll4* did not differ significantly between WT and *Mesp1*^Cre/+^/*Sox17*^fl/fl^embryos (Fig. 3*E*). At E8.5, the abundance of *Notch1* mRNA (Fig. 3*F*) and NOTCH1 protein (Fig. S7*G, H*) was unchanged in the mutant. However, at E9.5, the expression of *Notch1* and *Nrg1* (a downstream target of the NOTCH signal) in the heart was down-regulated significantly in the mutant embryos (Fig. 3*F, G*), suggesting that SOX17 does not directly induce NOTCH signaling but instead maintains it in the embryonic heart. The down-regulation of NOTCH signaling apparent at E9.5 is consistent with the previous finding that such signaling is required for trabeculation (37).

### Transcriptome Changes in Differentiating Cardiac Cells Induced by *Sox17* Deletion

The heart anomalies of mesoderm-specific *Sox17* mutant embryos were not apparent until E9.5, even though SOX17 is expected to function from E7.5 to E8.5 (Fig. 1; Figs. S1-S3). We therefore examined whether the loss of function for *Sox17* affects the endocardium from the earliest phase of its development, before the morphological abnormality is evident, with the use of single-cell microarray—based expression profiling of endocardium cells at E8.5 (nine-somite stage). Marked differences in gene expression profiles were detected between WT and *Mesp1*^Cre/+^/*Sox17*^fl/fl^cells. The normalized signal intensities of 114 and 171 probe sets (2.71% of total probe sets called “present”) were increased and decreased, respectively, by a factor of at least 5 in endocardium cells of the mutant relative to those of WT embryos (Fig. 4*A*; Table S1; *SI*, Gene Lists 1, 2). The gene expression profile of cardiomyocytes of E8.5 mutant embryos also differed from that of control cardiomyocytes, with the normalized signal intensities of 84 and 274 probe sets (4.78% of total probe sets called “present”) being increased and decreased, respectively (Fig. 4*B*; Table S1; *SI*, Gene Lists 3,4), supporting the notion that *Sox17* expression in the endocardium contributes to regulation of the genetic program for cardiomyocyte differentiation from the early stage of heart development.

**Fig. 4.**
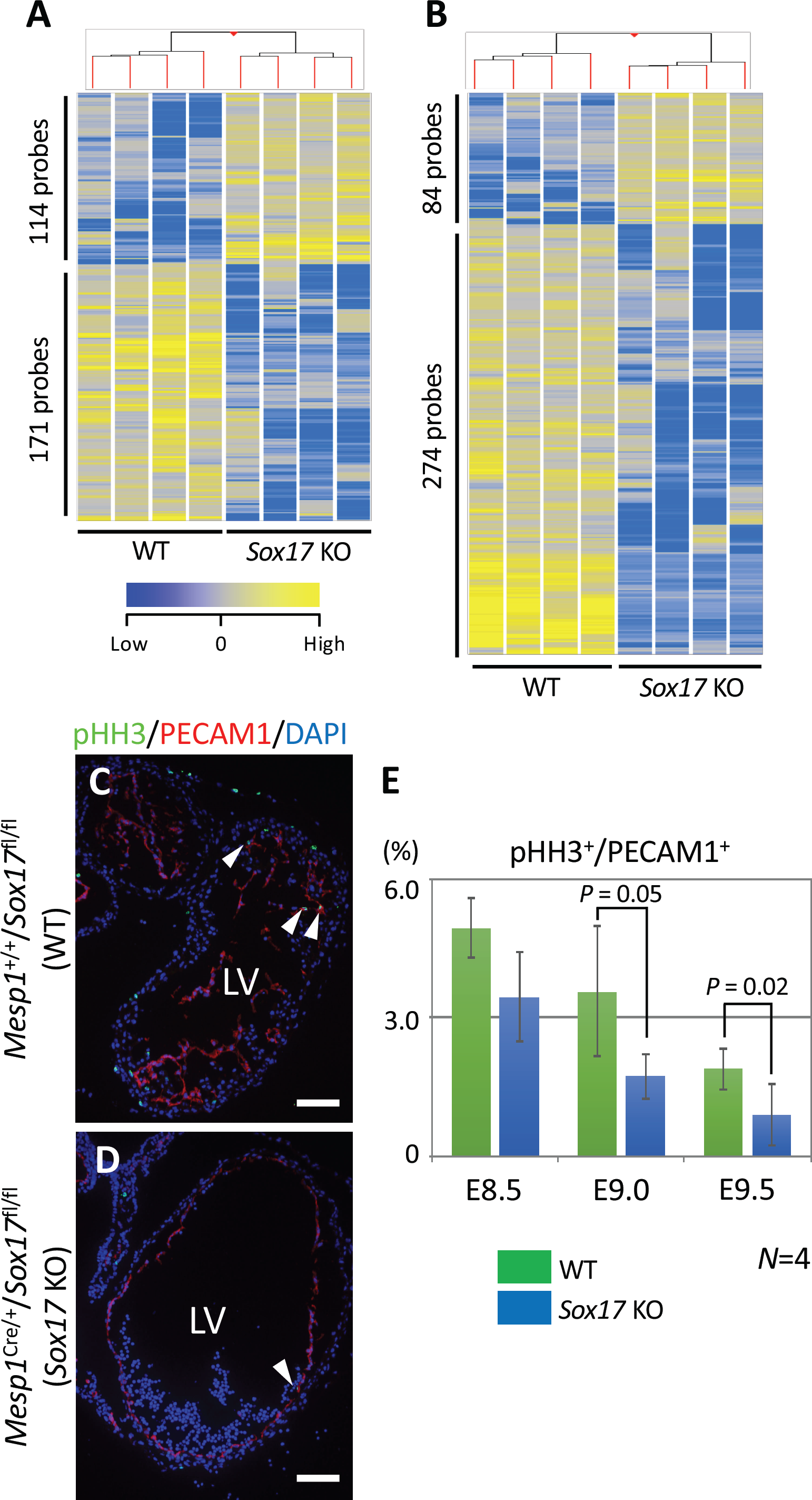
Altered gene expression profiles in the endocardium and myocardial cells of mesoderm-specific *Sox17* mutant embryos. (*A* and *B*) Hierarchical clustering and heat map for 285 (*A*) and 358 (*B*) probe sets for differentially expressed genes in the endocardium and myocardial cells, respectively, of WT and *Sox17* KO embryos at E8.5. (*C* and *D*) Immunofluorescence micrographs for pHH3 (green) and PECAM1 (red) in the heart of WT (*C*) and KO (*D*) embryos at E9.5. Blue is DAPI. Arrowheads indicate pHH3^+^endocardium cells. LV, left ventricle; Scale bars, 50 µm. (*E*) Proportion of pHH3^+^cells among PECAM1^+^cells in the heart of WT and *Sox17* KO embryos from E8.5 to E9.5. Means ± SD. Only *P* values of ≤0.05 (Student’s *t* test) are indicated.

Pathway analysis for the lists of genes whose expression was affected in mutant embryos revealed that molecules related to “cellular growth and proliferation,” “cellular development,” and “cell cycle” were significantly enriched for both endocardium cells and cardiomyocytes (Tables S2, S3). Examination of cell proliferation in the heart from E8.5 to E9.5 revealed that the ratio of KI67^+^(mitotic) endocardium cells or cardiomyocytes did not differ significantly between WT and mutant embryos at E8.5 or E9.0 but was significantly reduced in the mutant at E9.5. (Fig. S8*A, B*). Similarly, the ratio of phosphorylated histone H3 (pHH3)–positive (M-phase) cells in the endocardium of mutant embryos was significantly reduced at E9.0 (Fig. 4*C*–*E*). A significant reduction in the ratio of cardiomyocytes in M phase was apparent in the mutant at E9.5 (Fig. S8*C*–*E*), consistent with the notion that an initial anomalous molecular event in the endocardium subsequently affects the myocardium. Given that a reduction in *Notch1* expression in the mutant heart was not apparent until E9.5 (Fig. 3*F*), a primary event caused by the lack of *Sox17* expression in the endocardium might secondarily induce down-regulation of *Notch1* expression and thereby affect cell proliferation and trabeculation in the myocardium. Together, these results suggest that *Sox17* expression in the endocardium precursor cells is essential for regulation of endocardium development in a cell-autonomous manner and for that of myocardium development in a non–cell-autonomous manner.

## Discussion

We have here shown that *Sox17* is transiently expressed from E7.5 to E8.5 specifically in CPCs undergoing differentiation into the endocardium in mouse embryos. Gain—of-function and loss-of-function analyses revealed that *Sox17* is neither necessary nor sufficient for the induction of endocardial cell fate in CPCs. However, its expression in endocardium precursor cells is required for proper heart development and biases CPCs toward an endothelium-like phenotype. Our results suggest that SOX17 renders endocardium cells competent for proliferation and interaction with cardiomyocytes, with such interaction regulating cardiomyocyte growth and maturation (Fig. S8*F*).

Much of the available information relevant to endocardium differentiation has been derived from studies of the development of hematopoietic stem cells and the arterial endothelium. In mice, *Etv2* is essential for establishment of the endothelial and hematopoietic cell lineages in *Mesp1*^+^mesoderm, serving as a downstream target of bone morphogenetic protein (BMP), NOTCH, and WNT signaling pathways (12). Null mutation of *Etv2* results in embryonic mortality and loss of endothelial cells including the endocardium as well as of blood cells (12, 13). In addition, *Notch1*, which plays an important role in development of hemogenic arterial endothelial cells, is also required for development of the endocardium (38–40). Mesodermal cells expressing *Notch1* at E6.5 were found to contribute mostly to the endocardium (41). Given that *Sox17* appears to function in the hematopoietic lineage and that the endocardium likely possesses hemogenic ability (19, 42), the endocardium may share a common or similar developmental gene program including *Sox17* with the hematopoietic lineage. This notion is further supported by the observation that zebrafish mutants of *cloche*, which encodes a bHLH-PAS–type transcription factor and is expressed in endothelial and hematopoietic precursors, do not develop an endocardium or hematopoietic stem cells (43, 44). Of interest, Nkx2-5 was shown to transactivate *Etv2* in *Nkx2-5*^+^cells that contribute to the endocardium, suggesting that the cardiac program functions upstream of the endothelial-hemogenic progenitor gene program essential for endocardium development (45). However, the endothelial-hemogenic program including *Etv2* and *Tal1* (also known as *Scl*) prevents the induction of cardiomyocyte fate in CPCs (45, 46). Endocardium fate determination might thus be initiated by the cardiac program coupled to the endothelial-hemogenic program among *Mesp1*-expressing nascent mesoderm cells, with the endothelial-hemogenic program subsequently suppressing the cardiomyocyte program after commitment to the endocardium. Validation of the mechanism underlying fate determination for the endocardium will require further studies of the roles of *Mesp1, Sox17, Etv2, Notch1*, and other early endothelial-hemogenic factor genes.

*Sox17* has been shown to be required for vascular development. Its expression was thus found to promote sprouting angiogenesis during retinal vascularization and tumor angiogenesis (20, 47). Loss or gain of function of *Sox17* in the endothelial lineage of mouse embryos resulted in decreased and increased vascular density, respectively (48). SOX17 was suggested to destabilize vascular endothelial cells and thereby to secure the motility of tip cells during angiogenesis by regulating the proliferation, adhesion, and cytoskeletal organization of endothelial cells as well as the extracellular matrix. The phenotypes resulting from loss or gain of function of *Sox17* in our study are likely consistent with such a role, given the observed defects in cell shape, cell proliferation, and cell-cell interaction (Figs. 3, 4, Fig. S8). Endocardial cells that sprout toward the myocardium via an angiogenesis-like process may thus generate the touchdown sites at which cardiomyocytes are able to initiate their trabeculation (34, 35). For the endocardium to exert such an action, *Sox17* must be expressed at an appropriate level, given the phenotypes associated with its gain or loss of function (Figs. 2, 3). On the other hand, a specific role for *Sox17* in heart development is difficult to discern because of the functional redundancy of SOXF transcription factor genes. *Sox7* is expressed in a subset of the *Mesp1*^+^cell lineage from the earliest phase (41), and *Sox7* mutant embryos manifest an overall phenotype similar to that of embryos lacking *Sox17* (49). Identification of the direct targets of SOX17 and the other SOXF transcription factors should provide insight into the roles of these proteins in development.

Activated DLL4-NOTCH1 signaling in the endocardium was shown to be essential for ventricular trabeculation, with the consequent up-regulation of *Ephrinb2-EphB4-Nrg1* signaling promoting the differentiation of cardiomyocytes (35, 36). We found that loss of function of *Sox17* resulted in defective trabeculation, likely as a result of the significant down-regulation of *Notch1* and *Nrg1* expression in the heart apparent at E9.5. These findings appear consistent with the previous observation that SOX17 directly activates *Notch1* to establish the hemogenic endothelium (39).However, expression of *Notch1* was not affected in our *Sox17* loss-of-function mutant at E8.5, indicating that *Sox17* is dispensable for the induction of *Notch1* expression at the earliest phase of endocardium development, in contrast to the situation for hematopoietic stem cell development. The reduction in the level of *Notch1* expression in our *Sox17* mutant at E9.5 might be a secondary impairment due to a primary defect in the endocardium, or *Sox17* may be required for maintenance of *Notch1* expression. Alternatively, rather than SOX17, one or both of the other SOXF transcription factors (SOX7, SOX18) might directly activate *Notch1* in the endocardium lineage. Further studies of the relative roles of SOX17 and the other SOXF proteins should provide insight into the development of the endocardium.

## Materials and Methods

### BAC Transgenesis in mice

The BAC transgenesis was performed as previously described (23). All animal procedures were performed under project licenses (70/7254 and 70/7449) approved by the Home Office according to the Animals (Scientific Procedures) Act 1986 in the U.K., or with approval of the Osaka University Animal Experimentation Committee (license number: 29-039-004) in Japan.

### Real-Time PCR Analysis

Real-time PCR analysis with specific primer sets (*SI Appendix*, Table S4) was performed as previously described (5).

### Histology

Histological analyses were performed as previously described (5,23). The used primary and secondary antibodies were indicated in *SI Appendix*, Table S5.

### Single-cell microarray analysis

Single-cell cDNAs, constructed as previously described (23), were subjected to a SurePrint G3 Mouse GE v2 8×60K Microarray (Agilent). The microarray data obtained for this study are deposited in the Gene Expression Omnibus database (GEO, http://www.ncbi.nlm.nih.gov/geo) under the accession number GSE125323.

## ACKNOWLEDGMENTS

This work was supported by a U.K. Medical Research Council New Investigator Research Grant (G0900105) and a British Heart Foundation Project Grant (PG/11/102/29213) to K.Y.; by a Japan Society for the Promotion of Science KAKENHI Grant (17K08490) to R.S. We thank Nicholas Greene and Mitsuru Morimoto for technical advice as well as Brigid Hogan, Stefan Hoppler, Pete Scambler, and Paul Riley for helpful discussion.

## Author contributions

K.Y. designed research; K.Y., R.S., K.K., L.R., S.E., M.U., H.I., I.K., Y. Shintani, C.M., J.K., H.L., and K.S. performed research; K.Y., R.S., S.M., Y.K., M.A.-K., P.K., Y. Saga, and Y. Sawa analyzed data.; and K.Y. and R.S. wrote the paper.

The authors declare no conflict of interest.

## References

1. Lints TJ, et al (1993) Nkx-2.5?: a novel murine homeobox gene expressed in early heart progenitor cells and their myogenic descendants. Development 431:419–431.

2. Cai C, et al (2003) Isl1 Identifies a Cardiac Progenitor Population that Proliferates Prior to Differentiation and Contributes a Majority of Cells to the Heart. Dev Cell 5:877–889.

3. Bruneau BG, et al (2001) A Murine Model of Holt-Oram Syndrome Defines Roles of the T-Box Transcription Factor Tbx5 in Cardiogenesis and Disease. Cell 106:709–721.

4. Liang X, et al (2013) HCN4 Dynamically Marks the First Heart Field and Conduction System Precursors. Circ Res 113:399–407.

5. Ishida H, et al (2016) GFRA2 Identifies Cardiac Progenitors and Mediates Cardiomyocyte Differentiation in a RET-Independent Signaling Pathway. Cell Rep 16:1026–1038.

6. Kaufman M, Navaratnam V (1981) Early differentiation of the heart in mouse embryos. J Anat 133:235–246.

7. Deruiter MC, et al (1992) The development of the myocardium and endocardium in mouse embryos Fusion of two heart tubes? Anat Embryol (Berl) 185:461–473.

8. Misfeldt AM, et al (2009) Endocardial cells are a distinct endothelial lineage derived from Flk1 + multipotent cardiovascular progenitors. Dev Biol 333:78–89.

9. Lescroart F, et al (2014) Early lineage restriction in temporally distinct populations of Mesp1 progenitors during mammalian heart development. Nat Cell Biol 16:829–840.

10. Zamir L, et al (2017) Nkx2.5 marks angioblasts that contribute to hemogenic endothelium of the endocardium and dorsal aorta. eLlFE 6:1–31.

11. Bondue A, et al (2008) Mesp1 acts as a master regulator of multipotent cardiovascular progenitor specification. Cell Stem Cell 3:69–84.

12. Lee D, et al (2008) Article ER71 Acts Downstream of BMP, Notch, and Wnt Signaling in Blood and Vessel Progenitor Specification. Cell Stem Cell 2:497–507.

13. Ferdous A, et al (2009) Nkx2-5 transactivates the ETS-re;ated protein 71 gene and Specifies an Endothelial / Endocardial Fate in the Developing Embryo. Proc Natl Acad Sci 106:814–819.

14. Becker PW, et al (2016) An intronic Flk1 enhancer directs arterial-specific expression via RBPJ-mediated venous repression. Arterioscler Thromb Vasc Biol 36:1209–1219.

15. Wythe JD, Dang LTH, Devine WP, et al (2013) ETS factors regulate Vegf-dependent arterial specification. Dev Cell 26:45–58.

16. Shalaby F, et al (1995) Failure of blood-island formation and vasculogenesis in Dlk-1-deficient mice. Nature 376:62–66.

17. Gale NW, et al (2004) Haploinsufficiency of delta-like 4 ligand results in embryonic lethality due to majour deects in arterial and vascular development. Proc Natl Acad Sci 101:15949–15954.

18. Kanai-azuma M, et al (2002) Depletion of definitive gut endoderm in Sox17-null mutant mice. Development 129:2367–2379.

19. Kim I, Saunders TL, Morrison SJ (2007) Sox17 Dependence Distinguishes the Transcriptional Regulation of Fetal from Adult Hematopoietic Stem Cells. Cell 130:470–483.

20. Yang H, et al (2013) Sox17 promotes tumor angiogenesis and destabilizes tumor vessels in mice. J Clin Invest 123:418–431.

21. Engert S, et al (2009) Sox17-2A-iCre: A knock-in mouse line expressing Cre recombinase in endoderm and vascular endothelial cells. Genesis 47:603–610.

22. Liu Y, et al (2007) Sox17 is essential for the specification of cardiac mesoderm in embryonic stem cells. Proc Natl Acad Sci 104:3859–3864.

23. Kokkinopoulos I, et al (2015) Single-cell expression profiling reveals a dynamic state of cardiac precursor cells in the early mouse embryo. PLoS One 10:1–25.

24. Downs KM, Davies T (1993) Staging of gastrulating mouse embryos by morphological landmarks in the dissecting microscope. Development 118:1255–66.

25. Liao WP, et al (2009) Generation of a mouse line expressing Sox17-driven Cre recombinase with specific activity in arteries. Genesis 47:476–483.

26. Saga Y, et al (1999) MesP1 is expressed in the heart precursor cells and required for the formation of a single heart tube. Development 126:3437–47.

27. Srinivas S, Watanabe T, Lin C, et al (2001) Cre reporter strains produced by targeted insertion of EYFP and ECFP into the ROSA26 locus Shankar. BMC Dev Biol 1:4.

28. Rao PK, et al (2000) Isolation and Characterization of the Notch Ligand Delta4. Experimental Cell Res 260:379–386.

29. Baldwin HS, et al (1994) Platelet endothelial cell adhesion molecule-1 (PECAM-1/CD31): alternatively spliced, functionally distinct isoforms expressed during mammalian cardiovascular development. Development 120:2539–53.

30. Perou CM, et al (1997) Comparative mapping in the beige-satin region of mouse chromosome 13. Genomics 39:136–146.

31. Min B, Strauch AR, Foster DN (1988) Nucleic Acids Research Nucleotide sequence of a mouse vascular smooth muscle alpha-actin cDNA screened for vascular smooth muscle (VSM) alpha-actin cDNA probes for use in studies of smooth muscle myogenesis. Dideoxy chain termination sequence analysis. Nucleic Acids Res 16:10374.

32. Weinmaster G, Roberts VJ, Lemke G (1991) A homolog of Drosophila Notch expressed during mammalian development. Development 205:199–205.

33. Stanley EG, et al (2002) Efficient cre-mediated deletion in cardiac progenitor cells conferred by a 3’UTR-ires-Cre allele of the homeobox gene Nkx2-5. IntJDevBiol 46:431–439.

34. Jiménez-Amilburu V, et al (2016) In Vivo Visualization of Cardiomyocyte Apicobasal Polarity Reveals Epithelial to Mesenchymal-like Transition during Cardiac Trabeculation. Cell Rep 17:2687–2699.

35. Monte-Nieto G, et al (2018) Control of cardiac jelly dynamics by NOTCH1 and NRG1 defines the building plan for trabeculation. Nature. 557(7705):439–445.

36. Grego-Bessa J, et al (2007) Notch Signaling Is Essential for Ventricular Chamber Development. Dev Cell 12:415–429.

37. De la Pompa JL, Epstein JA (2012) Coordinating Tissue Interactions: Notch Signaling in Cardiac Development and Disease. Dev Cell 22:244–254.

38. Marcelo KL, et al (2013) Hemogenic endothelial cell specification requires c-Kit, notch signaling, and p27-mediated cell-cycle control. Dev Cell 27:504–515.

39. Clarke RL, et al (2013) The expression of Sox17 identifies and regulates haemogenic endothelium. Nat Cell Biol 15:502–510.

40. Richard C, et al (2013) Endothelio-Mesenchymal Interaction Controls runx1 Expression and Modulates the notch Pathway to Initiate Aortic Hematopoiesis. Dev Cell 24:600–611.

41. Lescroart F, et al (2018) Defining the earliest step of cardiovascular lineage segregation by single-cell RNA-seq. Science 359:1177–1181.

42. Nakano H, et al (2013) Haemogenic endocardium contributes to transient definitive haematopoiesis. Nat Commun 4:1–10.

43. Stainier DYR, et al (1995) cloche, an early acting zebrafish gene, is required by both the endothelial and hematopoietic lineages. Development 121:3141–3150.

44. Reischauer S, et al (2016) Cloche is a bHLH-PAS transcription factor that drives haemato-vascular specification. Nature 535:294–298.

45. Rasmussen TL, et al (2011) ER71 directs mesodermal fate decisions during embryogenesis. Development 138:4801–4812.

46. Van Handel B, et al (2012) Scl Represses Cardiomyogenesis in Prospective Hemogenic Endothelium and Endocardium. Cell 150:590–605.

47. Corada M, et al (2013) Sox17 is indispensable for acquisition and maintenance of arterial identity. Nat Commun 4:1–14.

48. Lee SH, et al (2014) Notch pathway targets proangiogenic regulator Sox17 to restrict angiogenesis. Circ Res 115:215–226.

49. Wat MJ, et al (2012) Mouse model reveals the role of SOX7 in the development of congenital diaphragmatic hernia associated with recurrent deletions of 8p23.1. Hum Mol Genet 21:4115–4125.

