## Supplementary Information for "*Sox17* expression in endocardium precursor cells regulates heart development in mice"

**SI Appendix**

Rie Saba *et al*.

**SI Inventory**

**Methods**

BAC Transgenesis

Mice

Staging of mouse embryos

RNA Isolation, RT, and Real-Time PCR Analysis

Histology

Single-cell cDNA Synthesis

**Tables**

Table S1. Microarray data and subtraction procedures.

Table S2. Pathway analysis for differentially expressed genes in endocardium cells of *Mesp1*$^{Cre/+}$/*Sox17*$^{fl/fl}$ versus *Mesp1*$^{+/+}$/*Sox17*$^{fl/fl}$ embryos at E8.5.

Table S3. Pathway analysis for differentially expressed genes in cardiomyocytes of *Mesp1*$^{Cre/+}$/*Sox17*$^{fl/fl}$ versus *Mesp1*$^{+/+}$/*Sox17*$^{fl/fl}$ embryos at E8.5.

Table S4. Primer sequences for real-time PCR analysis.

Table S5. Antibodies for immunofluorescence analysis.

**Figures**

Figure S1. SOX17 expression in the mesoderm cells of the mouse heart field.

Figure S2. Coexpression of SOX17 and NKX2-5 in a mouse embryo at the EHF stage (E7.5).

Figure S3. Distribution of SOX17$^{+}$ cells in the heart field of mouse embryos at the early somite stage.

Figure S4. Expression of arterial endothelial and cardiac marker genes in CPCs.

Figure S5. Mild to moderate gain-of-function phenotype for *Sox17* in mouse embryos.

Figure S6. Defective vascular remodeling in the yolk sac of mesoderm-specific *Sox17* mutant embryos at E9.5.

Figure S7. Anomalous looping of the heart tube in mesoderm-specific *Sox17* mutant embryos.

Figure S8. Reduced proliferation of cells in the endocardium and myocardium of mesoderm-specific *Sox17* mutant embryos.

**Gene lists**

Gene list 1. The 96 genes corresponding to the 114 probes in Results and Tables up-regulated in endocardial cells of *Mesp1*$^{Cre/+}$/*Sox17*$^{fl/fl}$ embryos relative to *Mesp1*$^{+/+}$/*Sox17*$^{fl/fl}$ embryos at E8.5.

Gene list 2. The 141 genes corresponding to the 171 probes down-regulated in endocardial cells of *Mesp1*$^{Cre/+}$/*Sox17*$^{fl/fl}$ embryos relative to *Mesp1*$^{+/+}$/*Sox17*$^{fl/fl}$ embryos at E8.5.

Gene list 3. The 64 genes corresponding to the 84 probes up-regulated in cardiomyocytes of *Mesp1*$^{Cre/+}$/*Sox17*$^{fl/fl}$ embryos relative to *Mesp1*$^{+/+}$/*Sox17*$^{fl/fl}$ embryos at E8.5.

Gene list 4. The 236 genes corresponding to the 274 probes down-regulated in cardiomyocytes of *Mesp1*$^{Cre/+}$/*Sox17*$^{fl/fl}$ embryos relative to *Mesp1*$^{+/+}$/*Sox17*$^{fl/fl}$ embryos at E8.5.

**References**

#### SI Methods

**BAC Transgenesis.** The construction of BAC transgene and transgenic mice production were performed as previously described (1, 2). All animal procedures were performed under project licenses (70/7254 and 70/7449) approved by the Home Office according to the Animals (Scientific Procedures) Act 1986 in the U.K., or with approval of the Osaka University Animal Experimentation Committee (license number: 29-039-004) in Japan. The BAC *Nkx2-5*$^{\mathrm{Sox17\text{-}IRES\text{-}LacZ\text{-}BghpA}}$ *Tg* was constructed with a BAC recombination system as shown in Figure 2*A*. The BAC clone RP24-324A10 contains 175.0 kb of the mouse *Nkx2-5* locus and drives expression of the *Sox17-IRES-LacZ* cassette (containing full-length mouse *Sox17* cDNA) according to the genomic context of *Nkx2-5*. A *Sox17-IRES-LacZ-BghpA* cassette was introduced in-frame into the mouse *Nkx2-5* gene of BAC clone RP24-324A10. For recombination with the BAC, left-arm XbaI-XhoI and right-arm EcoRI-EcoRV fragments were amplified by PCR independently with the primer sets *Nkx2-5*-L-arm-F (5′–XbaI site–GTCGACCGTTTAGACTCAGCATAACAG-3′) and *Nkx2-5*-L-arm-R (5′–XhoI site–CAGGTTTCACAGCGCCAGGTG-3′) as well as *Nkx2-5*-R-arm-F (5′–EcoRI site–GATAAAAAAGGTAAGGAGAAC-3′) and *Nkx2-5*-R-arm-R (5′-EcoRV site–GGCAGGGTGGGCTACACAAGG-3′), respectively. The right-arm EcoRI-EcoRV fragment was cloned into pL453 as *Frt–Neo*$^r$*–Frt*–R arm, and the left-arm XbaI-XhoI and *IRES-LacZ* XhoI-BamHI fragments were then simultaneously introduced to yield L arm–*IRES–LacZ–Frt–Neo*$^r$*–Frt*–R arm. A XhoI-BglII fragment of mouse *Sox17* cDNA obtained by PCR with the primers 5′–XhoI site–GTCGCCACCATGAGCAGCCCGGATGCGGGA-3′ and 5′-TCTGCGTTGTGCAGATCTGGG-3′ and a BglII-XbaI fragment of the *Sox17* cDNA were simultaneously cloned into the plasmid to yield L arm–*Sox17–IRES–LacZ–Frt–Neo*$^r$*–FRT*–R arm. EL250 cells transformed with the RP24-324A10 BAC clone were subjected to electroporation with the L arm–*Sox17–IRES–LacZ–Frt–Neo*$^r$*–FRT*–R arm fragment and then to selection with kanamycin. Following removal of the *Neo*$^r$ cassette by arabinose treatment (Flp induction), the BAC *Tg* was prepared and used for microinjection.

**Mice.** *Sox17*$^{\mathrm{fl}}$ (MGI ID: 3717121) (3) and *Rosa26*$^{\mathrm{EYFP}}$ Cre reporter (MGI ID: 2449038) (4) mice were obtained from The Jackson Laboratory. *Mesp1*$^{\mathrm{Cre}}$ mice (MGI ID:

2176467) were described previously (5).

**Staging of mouse embryos**

Developmental stages of mouse embryos were classified according to morphology as previously described (6). The morning of the day of vaginal plug detection was set as E0.5.

**RNA Isolation, RT, and Real-Time PCR Analysis.** Embryonic hearts were dissected in ice-cold phosphate-buffered saline (PBS). Total RNA was isolated with the use of an RNeasy Mini Kit (Qiagen) and was subjected to reverse transcription (RT) with the use of Superscript III (ThermoFisher Scientific) and an oligo(dT) primer. The resulting cDNA was subjected to real-time PCR analysis with specific primer sets (Table S4) and with the use of a QuantiTect SYBR Green PCR Kit and Rotorgene (Qiagen). Data were normalized by *Gapdh* expression.

**Histology.** Mouse embryos were dissected in ice-cold PBS and fixed with 4% paraformaldehyde in PBS at 4°C for 2 h. For assay of BrdU incorporation, pregnant mice were injected intraperitoneally with 500 μg of BrdU (Sigma-Aldrich) at 3 h before embryo removal. Fixed embryos were embedded in OCT compound (Sakura Finetek), frozen, sectioned at a thickness of 8 μm, and stained with primary and secondary antibodies (Table S5) as previously described (2, 7). DNA was counterstained with DAPI (Merck). Immunofluorescence micrographs were acquired with an LSM510 confocal (Zeiss) or BZ8000 fluorescence (Keyence) microscope. Section in situ hybridization was performed as described (2, 7).

**Single-cell microarray analysis.** The dissected hearts were treated with trypsin for 3 min at 37°C to isolate single cells. First-strand DNA was synthesized for 10 min at 50°C with Superscript III reverse transcriptase (ThermoFisher Scientific). The cell type for each single-cell cDNA preparation was identified by PCR of marker gene expression with specific primer sets (Table S4). The T3 promoter was added to the 5′ end of each cDNA by PCR with the T3V3 primer (5′-
CCAAGCTCGAAATTAACCCTCACTAAAGGGAGAATATCTCGAGGGCGCGCCG

GATCC-3′) and V1 $dT_{24}$ primer (5′-ATATGGATCCGGCGCGCCGTCGACTTTTTTTTTTTTTTTTTTTTTTTT-3′). Cells were isolated from the heart ventricle of control (*Sox17* $^{fl/fl}$) or *Sox17* knockout (*Mesp1*$^{Cre/+}$/*Sox1* $^{fl/fl}$) embryos at E8.5, and single-cell cDNA was synthesized by RT as previously described with some modifications (2). The T3 promoter was added to the 5′ end of each cDNA by PCR. Virtual mRNAs were synthesized from the T3 promoter–cDNA constructs with a MEGAScript T3 Transcription Kit (ThermoFisher Scientific), and they (50 ng) were then labeled with Cy3 with the use of a Low RNA Input QuickAmp (One Color) Labeling Kit before hybridization with a SurePrint G3 Mouse GE v2 8×60K Microarray with the use of a Gene Expression Hybridization Kit (Agilent). The processed data were analyzed statistically with Genespring GX software (Agilent). Expression levels of <50 were set to 50, per chip normalization was based on the 50th percentile, and per gene normalization was based on the median. Gene subtraction was performed as shown in Table 1. Gene lists were analyzed with Ingenuity Pathway Analysis software (Qiagen Bioinformatics). The microarray data obtained for this study are deposited in the Gene Expression Omnibus database (GEO, http://www.ncbi.nlm.nih.gov/geo) under the accession number GSE125323.

**SI Tables**

**Table S1.** Microarray data and subtraction procedures.

| Subtraction procedure | Endocardium<br>E8.5<br>(nine-somite) | Myocardium<br>E8.5<br>(nine-somite) |
|---|---|---|
| All probe sets | 55821 | 55821 |
| a) Flags detected in at least one condition | 10496 | 7485 |
| b) Expression level of >50 in at least one condition | 5329 | 4515 |
| c) *P* < 0.05 (*t* test) for WT vs. KO | 460 | 712 |
| d) Fivefold change for WT vs. KO | 285 | 358 |
| e) d/a (%) | 2.71 | 4.78 |
| Increased in KO | 114 | 84 |
| Decreased in KO | 171 | 274 |

Probe sets whose expression in endocardium or myocardium cells of embryos at E8.5 (nine-somite stage) was affected by knockout (KO) of *Sox17* in mesoderm were extracted by filtering steps: a) "present" either in *Mesp1*$^{+/+}$/*Sox17*$^{fl/fl}$ (WT) or *Mesp1*$^{Cre/+}$/*Sox17*$^{fl/fl}$ (KO) embryos; b) a raw expression value of >50; c) significant difference between WT and KO by Student's *t* test (*P* < 0.05); d) fivefold difference in normalized signals between WT and KO; e) calculation of d/a ratio.

**Table S2.** Pathway analysis for differentially expressed genes in endocardium cells of *Mesp1*$^{Cre/+}$/*Sox17*$^{fl/fl}$ versus *Mesp1*$^{+/+}$/*Sox17*$^{fl/fl}$ embryos at E8.5.

| Molecular and cellular functions | *P* value | No. of molecules |
|---|---|---|
| Cellular growth and proliferation | 1.29E-02 – 3.29E-08 | 90 |
| Cellular development | 1.31E-02 – 3.52E-06 | 65 |
| Cell cycle | 1.24E-02 – 8.97E-06 | 35 |
| DNA replication, recombination, and repair | 9.65E-03 – 2.94E-05 | 25 |
| Gene expression | 1.32E-02 – 3.30E-05 | 53 |

Enriched pathways in the category of "Molecular and cellular functions" were determined by Ingenuity Pathway Analysis for the 285 probe sets in *SI Appendix*, Gene Lists 1 and 2.

**Table S3.** Pathway analysis for differentially expressed genes in cardiomyocytes of *Mesp1*$^{Cre/+}$/*Sox17*$^{fl/fl}$ versus *Mesp1*$^{+/+}$/*Sox17*$^{fl/fl}$ embryos at E8.5.

| Molecular and cellular functions | *P* value | No. of molecules |
|---|---|---|
| Cell cycle | 2.27E-02 – 2.24E-09 | 43 |
| Cellular development | 2.27E-02 – 2.24E-09 | 69 |
| RNA posttranscriptional modification | 2.27E-02 – 5.04E-07 | 19 |
| Cellular growth and proliferation | 2.27E-02 – 9.13E-05 | 95 |
| Cellular assembly and organization | 2.27E-02 – 1.12E-04 | 17 |

Enriched pathways in the category of "Molecular and cellular functions" were determined by Ingenuity Pathway Analysis for the 358 probe sets in *SI Appendix*, Gene Lists 3 and 4.

**Table S4. Primer sequence for RT-PCR.**

| Gene name | Sense primer (5'-3') | Antisense primer (5'-3') |
|---|---|---|
| *Cryptic* | CGCCAGAGGATCAAGAGAAT | CTTAGGAGCCTCAGCCCTTA |
| *Nkx2-5* | TTGACGTAGCCTGGTGTCTC | TAGTGTGGAATCCGTCGAAA |
| *Tbx5* | CGTCGTGGAATTCAGAGTTG | CGATAGGTGCTGAGGAGTGA |
| *Sox17* | TTCTGTACACTTTAATGAGGCTGTTC | TTGTGGGAAGTGGGA TCAAG |
| *Sox2* | CATGAGAGCAAGTACTGGCAAG | CCAACGATATCAACCTGCATGG |
| *Dll4* | AGCTGTGACTCCTGCCTCCAACCCC | CACATGAGCCCAGAGGCACCAGGAC |
| *Pecam1* | CGGTTTCCTAAGGTCTGAGC | AGGCGAGGAGGGTTAGGTAT |
| *Actn2* | CAGGTCATCGCCTCCTTCCGGATT | GTCGCTTTCCCCGTAGAGGGCAGA |
| *Acta2* | TGTGTTTCTGTTAGGTGAGAATCA | TGGACCTTCCTCTGTTGAAGT |
| *Etv2* | AATAGCCGCGAGTTCCAG | CATAATTCATTCCCGGCTTC |
| *Notch1* | TCTGCTGCCCACCAAGTCCCTTTCC | TTAGGCATGGCACAGACACTGCCCC |
| *Nrg1* | GTATCAGCCATGACCCCCGGCTC | GCTGACTGCCACAGAGGGCATGGAC |
| *Gapdh* | ACAGTCCATGCCATCACTGCCACCC | CACAGCCTTGGCAGCACCAGTGGAT |

**Table S5. Antibodies for immunohistochemistry.**

| Primary Antibodies | Target protein | Immune Animal | Company | Product No. | Dilution |
|---|---|---|---|---|---|
| Rb mAb to GFP | GFP | Rabbit | abcam | ab32146 | 1:1000 |
| Anti-Sox17 | Sox17 | Goat | R&D | AF1924 | 1:100 |
| Goat anti-cTNT | cardiac Troponin T | Goat | Hytest | 4T19/2 | 1:3000 |
| Rat anti-CD31 (MEC 13.3) | CD31 | Rat | B&D Bioscience | 550274 | 1:500 |
| Ms mAb to MHC (MF20) | Myosin Heavy Chain | Mouse | Novus Biologicals | MAB4470 | 1:1000 |
| Cleaved Caspase-3 (D175) Rabbit Ab | Cleaved Caspase-3 | Rabbit | CST | 9661S | 1:100 |
| Notch1 (D1E11) XP(R) Rabbit mAb | Notch1 | Rabbit | CST | 3608S | 1:100 |
| Rb pAb to Ki67 | Ki67 | Rabbit | abcam | ab15580 | 1:100 |
| Phospho-Histone H3 (Ser10) antibody | p-H3 | Rabbit | CST | 9708 | 1:100 |
| Rb pAB Nkx2-5 | Nkx2-5 | Rabbit | abcam | ab35842 | 1:100 |
| Histone H3 (D1H2) XP(R) Rabbit mAb | Histone H3 | Rabbit | CST | 4499S | 1:100 |
| Mouse anti-BrdU (BU33) | BrdU | Mouse | Sigma | BU33 | 1:1000 |
| Secondary Antibodies | Target protein | Immune Animal | Company | Product No. | Dulution |
| Donkey anti-rabbit IgG Alexa 488 | rabbit IgG | Donkey | Jackson ImmunoResearch Laboratories | 711-546-152 | 1:1000 |
| Donkey anti-rat IgG Alexa 594 | rat IgG | Donkey | Jackson ImmunoResearch Laboratories | 712-586-153 | 1:1000 |
| Donkey anti-goat IgG Alexa 488 | goat IgG | Donkey | Jackson ImmunoResearch Laboratories | 705-546-147 | 1:1000 |
| Donkey anti-mouse IgG Alexa 594 | mouse IgG | Donkey | Jackson ImmunoResearch Laboratories | 715-586-151 | 1:1000 |
| Donkey anti-mouse IgG Alexa 488 | mouse IgG | Donkey | Jackson ImmunoResearch Laboratories | 715-546-151 | 1:1000 |

**SI Figure Legends**

**Fig. S1**. SOX17 expression in the mesoderm of the mouse heart field. (*A*) Schematic representation of a mouse embryo at the early head fold stage (E7.5) as a left lateral view. The magenta line shows the sectional plane along the anterior (A)–posterior (P) axis in *B*. (*B* and *C*) Immunofluorescence micrographs of SOX17$^{+}$ cells (red) in a *Mesp1*$^{Cre/+}$/*Rosa26*$^{EYFP/+}$ mouse embryo. EYFP (green) marks mesoderm cells, and nuclei (blue) were stained with 4′,6-diamidino-2-phenylindole (DAPI). The boxed region in *B* is shown at higher magnification in *C*. Arrowheads indicate EYFP$^{+}$, SOX17$^{+}$ cells. Al, allantoic bud. Scale bar, 100 μm.

**Fig. S2.** Coexpression of SOX17 and NKX2-5 in a mouse embryo at the EHF stage (E7.5). Expression of SOX17 (green) and NKX2-5 (red) was examined by immunofluorescence analysis. Nuclei (blue) were stained with DAPI. The boxed regions in *A* are shown at higher magnification in *B* and *C*. Arrows and arrowheads indicate NKX2-5 single-positive cells and cells positive for both SOX17 and NKX2-5, respectively. Al, allantois. Scale bars, 100 μm (*A*) and 50 μm (*B* and *C*).

**Fig. S3.** Distribution of SOX17$^{+}$ cells in the heart field of mouse embryos at the early somite stage. Expression of SOX17 (red) and EYFP (green) in the heart field of *Mesp1*$^{Cre/+}$/*Rosa26*$^{EYFP/+}$ mouse embryos at the zero-somite (*A*) and three-somite (*B*) stages was examined by immunofluorescence analysis. Nuclei (blue) were stained with DAPI. The boxed region in the upper left panel for each embryo is shown at higher magnification in the corresponding panels labeled i to iii. Arrowheads indicate cells positive for both SOX17 and EYFP. An, anterior; Ps, posterior. Scale bars, 100 μm.

**Fig. S4.** Expression of arterial endothelial and cardiac marker genes in CPCs. The ratio of *Dll4*$^{+}$ or *Pecam1*$^{+}$ cells (*A*), *Actn2*$^{+}$ or *Acta2*$^{+}$ cells (*B*) or of *Etv2*$^{+}$ or *Notch1*$^{+}$ cells (*C*) among various categories of CPCs in mouse embryos at the indcated stages was determined by PCR analysis of single-cell cDNA. The number of cells analyzed is shown in parentheses.

**Fig. S5.** Mild to moderate gain-of-function phenotype for *Sox17* in mouse embryos. (*A* and *B*) Fluorescence micrographs for immunostaining of MYH1 (cardiomyocytes, green and isolectin B4 staining (endocardium, red) in the heart of WT (*A*) and BAC *Nkx2-*

$5^{Sox17\text{-}IRES\text{-}LacZ\text{-}BghA}$ transgenic (*B*) embryos at E9.5. Nuclei (blue) were stained with DAPI. Arrowheads indicate aggregated endocardium cells. Scale bars, 20 µm. (*C–E*) Immunofluorescence micrographs for MYH1 (red) and cleaved caspase-3 (apoptotic cells, green) in *Tg*(–) (*C*) and *Tg*(+) (*D*) embryos at E9.5. Nuclei (blue) were stained with DAPI. The boxed region in *D* is shown at higher magnification in *E*. The arrow and arrowheads indicate apoptotic cells positive for cleaved caspase-3 in the endocardium and myocardium, respectively Scale bar, 100 µm. (*F* and *G*) Immunofluorescence micrographs for bromodeoxyuridine (BrdU) incorporation (red) as well as for NKX2-5 (green) and histone H3 (HH3, nucleus, blue) in *Tg*(–) (*F*) and *Tg*(+) (*G*) embryos at E8.. Scale bar, 100 µm. (*H*) Proportions of $BrdU^{+}$ cells among $NKX2\text{-}5^{+}$ cells in the heart of *Tg*(–) and *Tg*(+) embryos at E8.5. Data are means ± SD ($N = 3$ embryos). The *P* value was determined by Student's *t* test.

**Fig. S6.** Defective vascular remodeling in the yolk sac of mesoderm-specific *Sox17* mutant embryos at E9.5. (*A* and *B*) Macroscopic observation of the yolk sac of $Mesp1^{+/+}/Sox17^{fl/fl}$ (*A*) and $Mesp1^{Cre/+}/Sox17^{fl/fl}$ (*B*) embryos. Remodeled large blood vessels similar to those indicated by the arrowheads in *A* are not apparent in the mutant. Scale bar, 500 µm. (*C* and *D*) Giemsa staining of the yolk sac of WT (*C*) and mutant (*D*) embryos. Blood islands (arrows) containing primary erythrocytes are formed in both WT and mutant embryos, whereas remodeled blood vessels (arrowheads) are present only in the WT embryo. Upper and lower panels show different areas of the yolk sac. Scale bar, 100 µm.

**Fig. S7.** Anomalous looping of the heart tube in mesoderm-specific *Sox17* mutant embryos. (*A–F*) The heart of $Mesp1^{+/+}/Sox17^{fl/fl}$ (*A–C*) or $Mesp1^{Cre/+}/Sox17^{fl/fl}$ (*D–F*) embryos at the 22-somite stage (E9.5) is shown in right lateral (*A* and *D*), frontal (*B* and *E*), and left lateral (*C* and *F*) views. Arrows and arrowheads indicate the borders of the outflow tract (OFT) and ventricle (V) and of the ventricle and atrium (A), respectively. (*G* and *H*) Immunofluorescence micrographs for NOTCH1 (green) and TNNT (red) in the heart of $Mesp1^{+/+}/Sox17^{fl/fl}$ (*G*) and $Mesp1^{Cre/+}/Sox17^{fl/fl}$ (*H*) embryos at the nine-somite stage (E8.5). Nuclei (blue) were stained with DAPI. Ec, endocardium; Mc, myocardium. Scale bar, 100 µm.

**Fig. S8.** Reduced proliferation of cells in the endocardium and myocardium of mesoderm-specific *Sox17* mutant embryos. (*A* and *B*) Proportions of $KI67^{+}$ cells among $PECAM1^{+}$ endocardium cells (*A*) or $TNNT^{+}$ cardiomyocytes (*B*) of *Mesp1*$^{+/+}$/*Sox17*$^{fl/fl}$ (control) and *Mesp1*$^{Cre/+}$/*Sox17*$^{fl/fl}$ (*Sox17* KO) embryos from E8.5 to E9.5. Data are means ± SD ($N = 4$ embryos). Only significant $P$ values calculated by Student's $t$ test are shown. (*C* and *D*) Immunofluorescence micrographs of pHH3 (green) and TNNT (red) in the myocardium of the left ventricle (LV) of *Mesp1*$^{+/+}$/*Sox17*$^{fl/fl}$ and *Mesp1*$^{Cre/+}$/*Sox17*$^{fl/fl}$ embryos at E9.5. Nuclei (blue) were stained with DAPI. Arrowheads indicate pHH3-positive cells. Scale bars, 50 µm. (*E*) Proportion of $pHH3^{+}$ cells among $TNNT^{+}$ cardiomyocytes in mutant and control embryos from E8.5 to E9.5. Data are means ± SD ($N = 4$ embryos). Only significant $P$ values calculated by Student's $t$ test are shown. (*F*) Model for the role of *Sox17* in endocardium development and function. ECPC, endocardium precursor cell.

#### *SI* References


1. Uehara M, et al (2009) Removal of maternal retinoic acid by embryonic CYP26 is required for correct Nodal expression during early embryonic patterning. Genes Dev 23:1689–1698.
2. Kokkinopoulos I, et al (2015) Single-cell expression profiling reveals a dynamic state of cardiac precursor cells in the early mouse embryo. PLoS One 10:1–25.
3. Kim I, Saunders TL, Morrison SJ (2007) Sox17 Dependence Distinguishes the Transcriptional Regulation of Fetal from Adult Hematopoietic Stem Cells. Cell 130:470–483.
4. Srinivas S, et al (2001) Cre reporter strains produced by targeted insertion of. BMC Dev Biol 1:4.
5. Saga Y, et al (1999) MesP1 is expressed in the heart precursor cells and required for the formation of a single heart tube. Development 126:3437–47.
6. Downs KM, and Davies T. (1993). Staging of gastrulating mouse embryos by morphological landmarks in the dissecting microscope. Development 118, 1255-1266.
7. Ishida H, et al (2016) GFRA2 Identifies Cardiac Progenitors and Mediates Cardiomyocyte Differentiation in a RET-Independent Signaling Pathway Article GFRA2 Identifies Cardiac Progenitors and Mediates Cardiomyocyte Differentiation in a RET-Independent Signaling Pathway. Cell Rep 16:1026–1038.

Figure S1

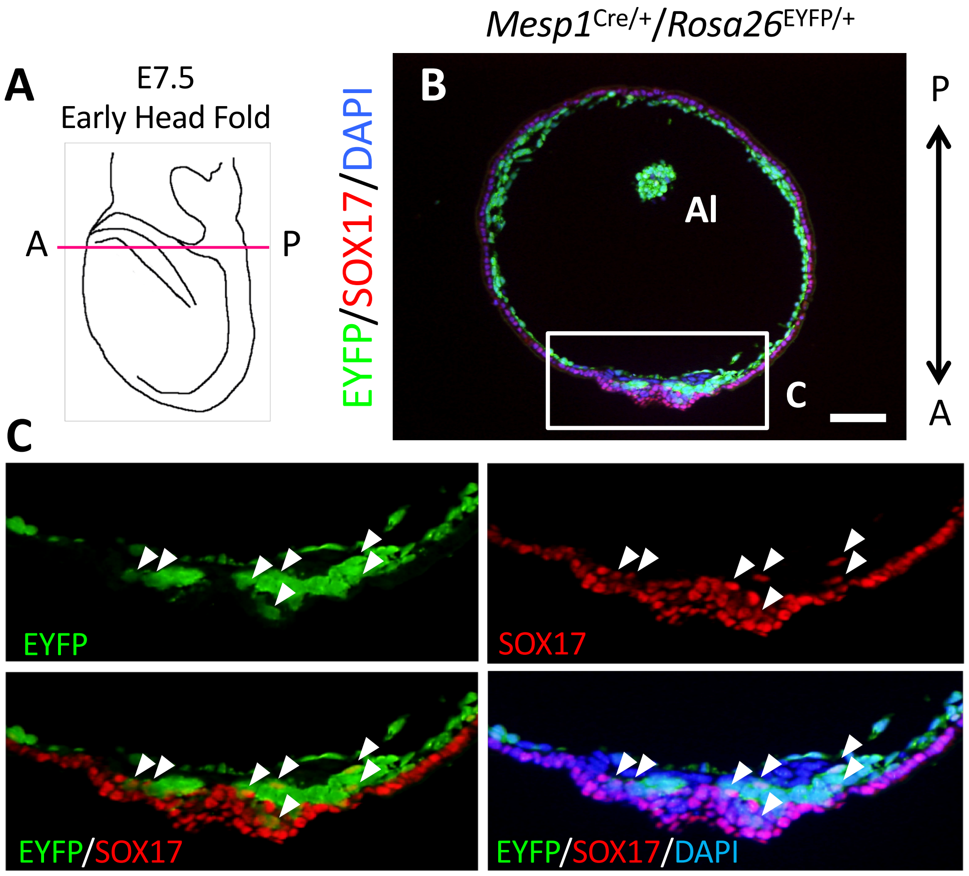

A
E7.5
Early Head Fold
A
P
*Mesp1*Cre/+/*Rosa26*EYFP/+
EYFP/SOX17/DAPI
B
Al
C
P
A
C
EYFP
SOX17
EYFP/SOX17
EYFP/SOX17/DAPI

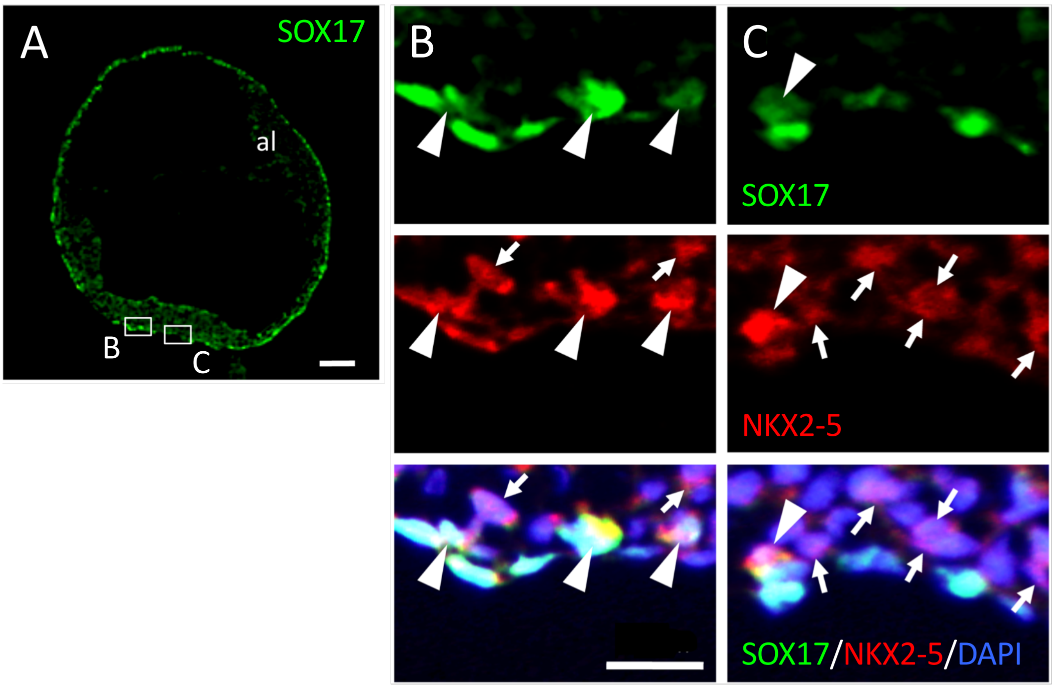
A
SOX17
al
B
C
B
C
SOX17
NKX2-5
SOX17/NKX2-5/DAPI

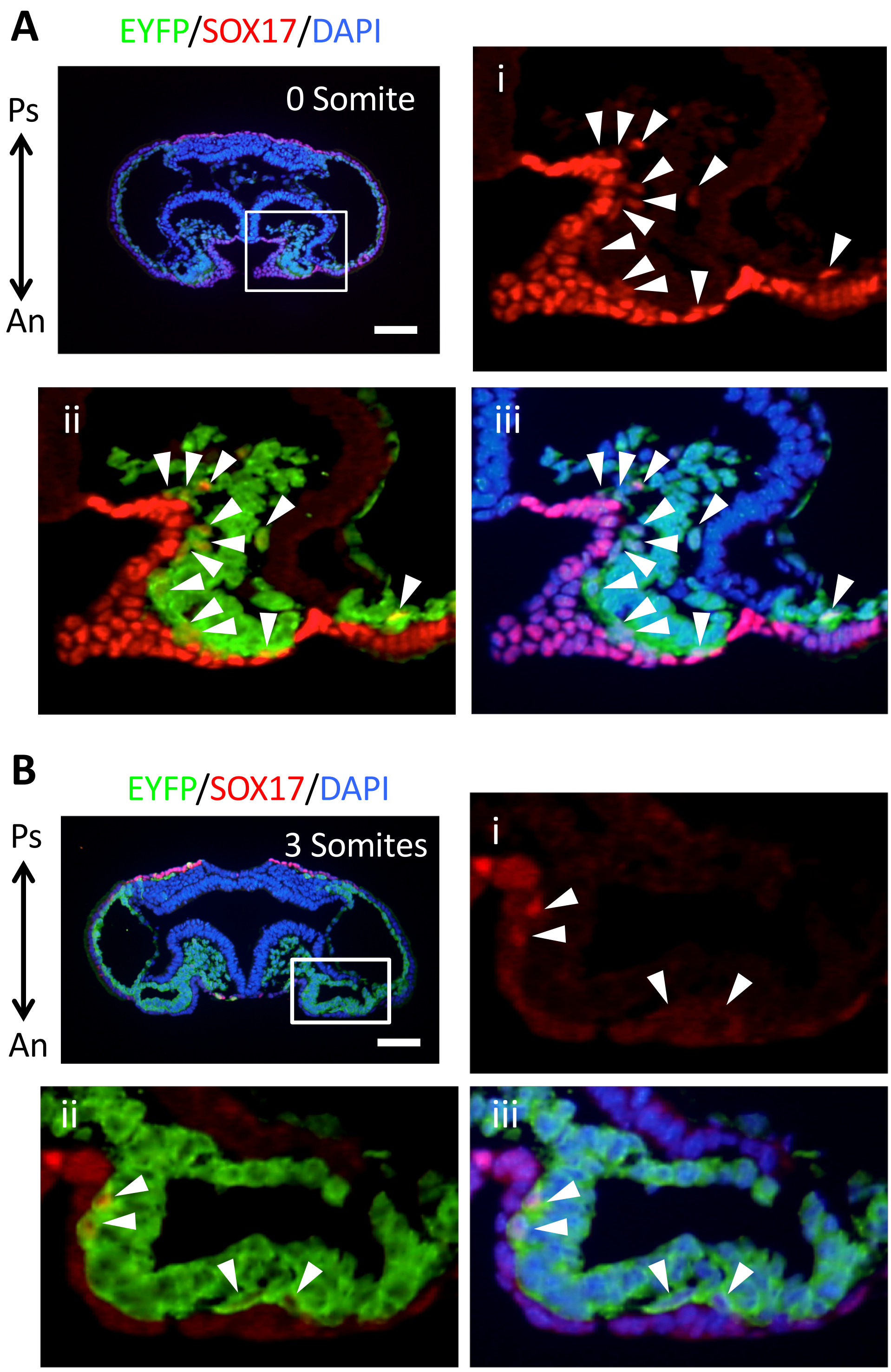
A
EYFP/SOX17/DAPI
Ps
An
0 Somite
i
ii
iii
B
EYFP/SOX17/DAPI
Ps
An
3 Somites
i
ii
iii

### Figure S4

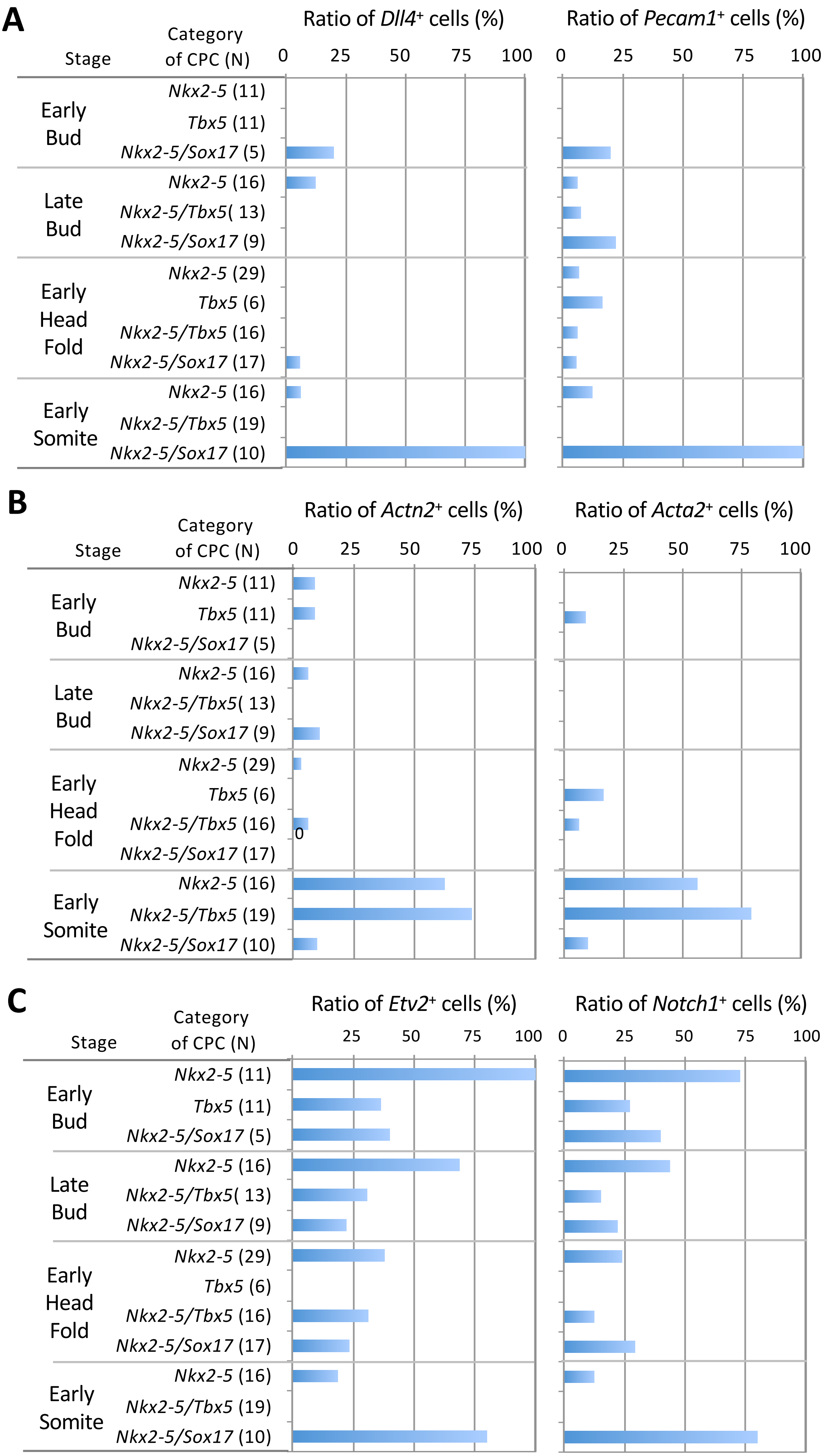


Saba et al., 2019, Supplement

### Figure S5

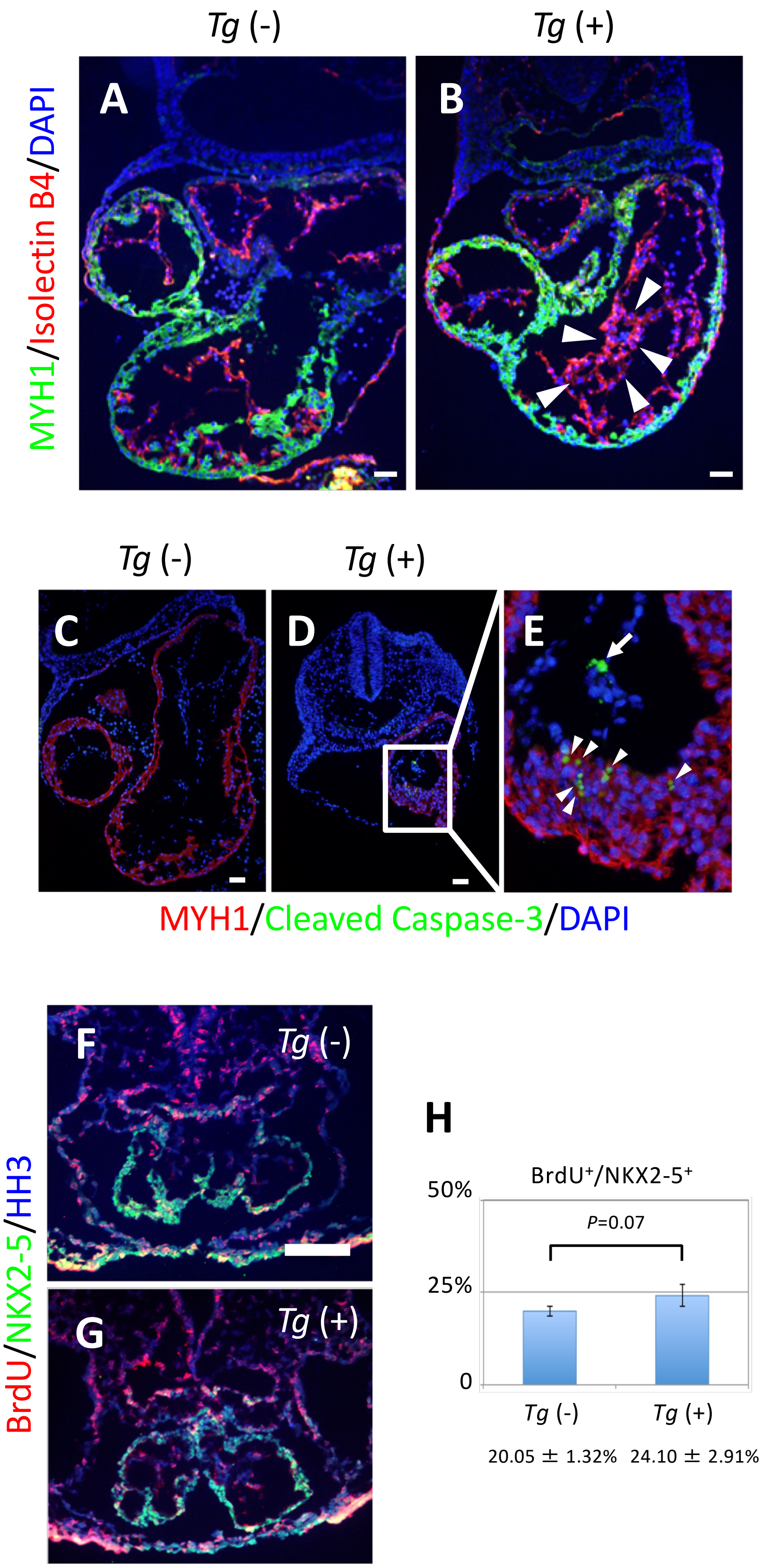

Tg (-)
Tg (+)
A
B
MYH1/Isolectin B4/DAPI
Tg (-)
Tg (+)
C
D
E
MYH1/Cleaved Caspase-3/DAPI
F
Tg (-)
G
Tg (+)
BrdU/NKX2-5/HH3
H
BrdU+/NKX2-5+
50%
25%
0
P=0.07
Tg (-)
Tg (+)
20.05 ± 1.32%
24.10 ± 2.91%

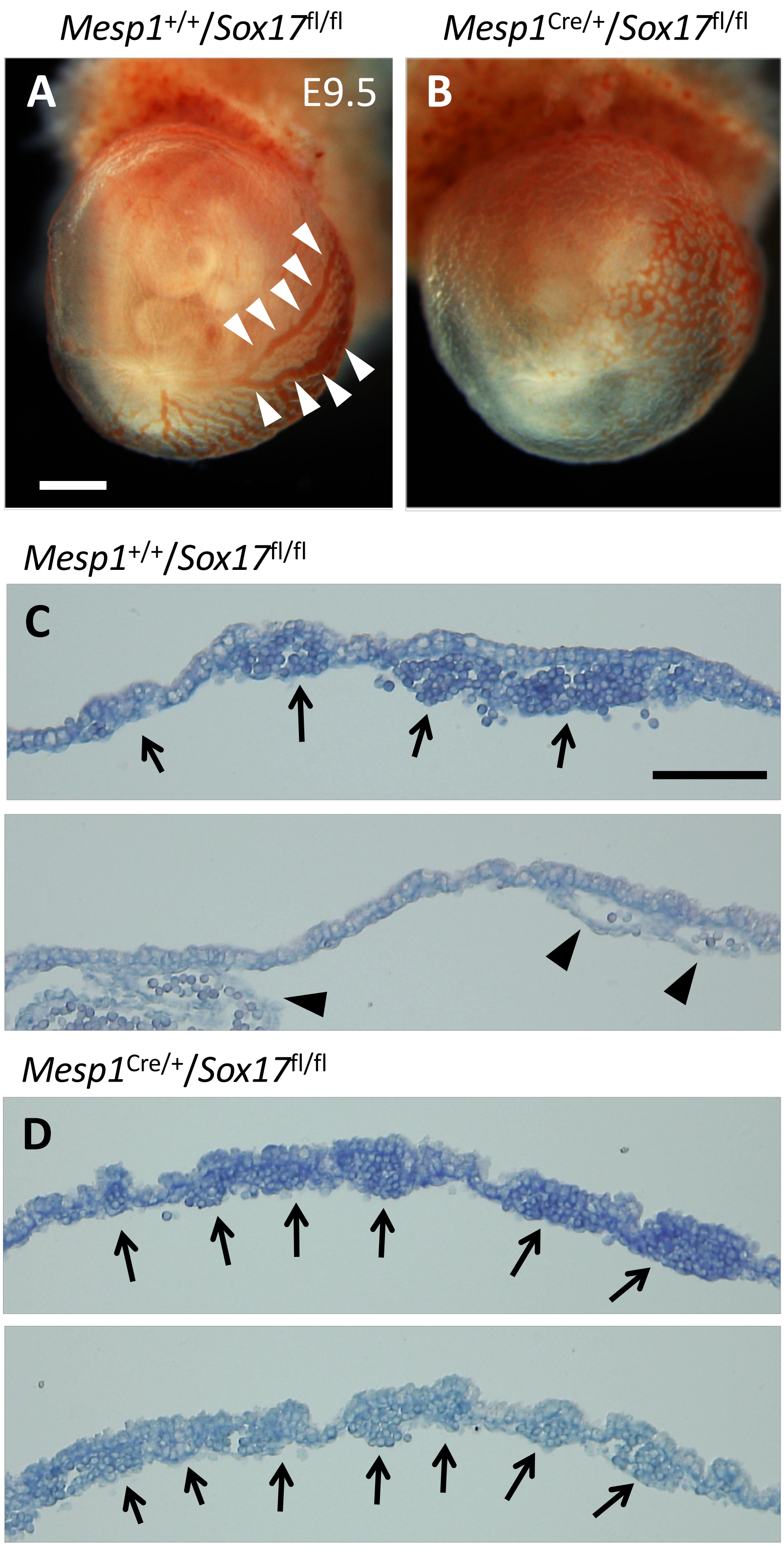
Mesp1+/+/Sox17fl/fl
Mesp1Cre/+/Sox17fl/fl
A
E9.5
B
Mesp1+/+/Sox17fl/fl
C
Mesp1Cre/+/Sox17fl/fl
D

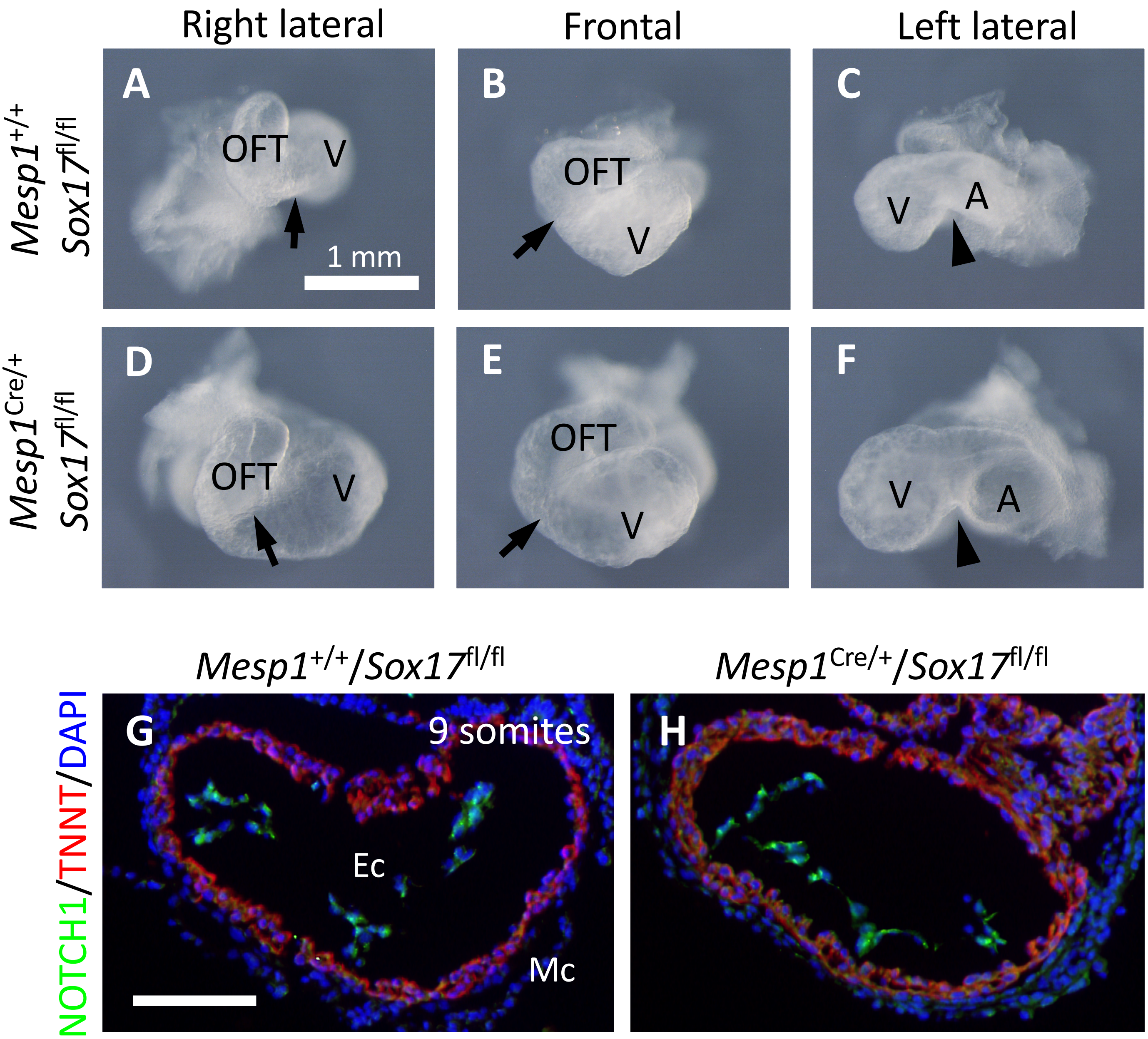
Right lateral
Frontal
Left lateral
A
B
C
D
E
F
OFT
V
A
1 mm
Mesp1+/+ Sox17fl/fl
Mesp1Cre/+ Sox17fl/fl
Mesp1+/+/Sox17fl/fl
Mesp1Cre/+/Sox17fl/fl
NOTCH1/TNNT/DAPI
G
H
9 somites
Ec
Mc

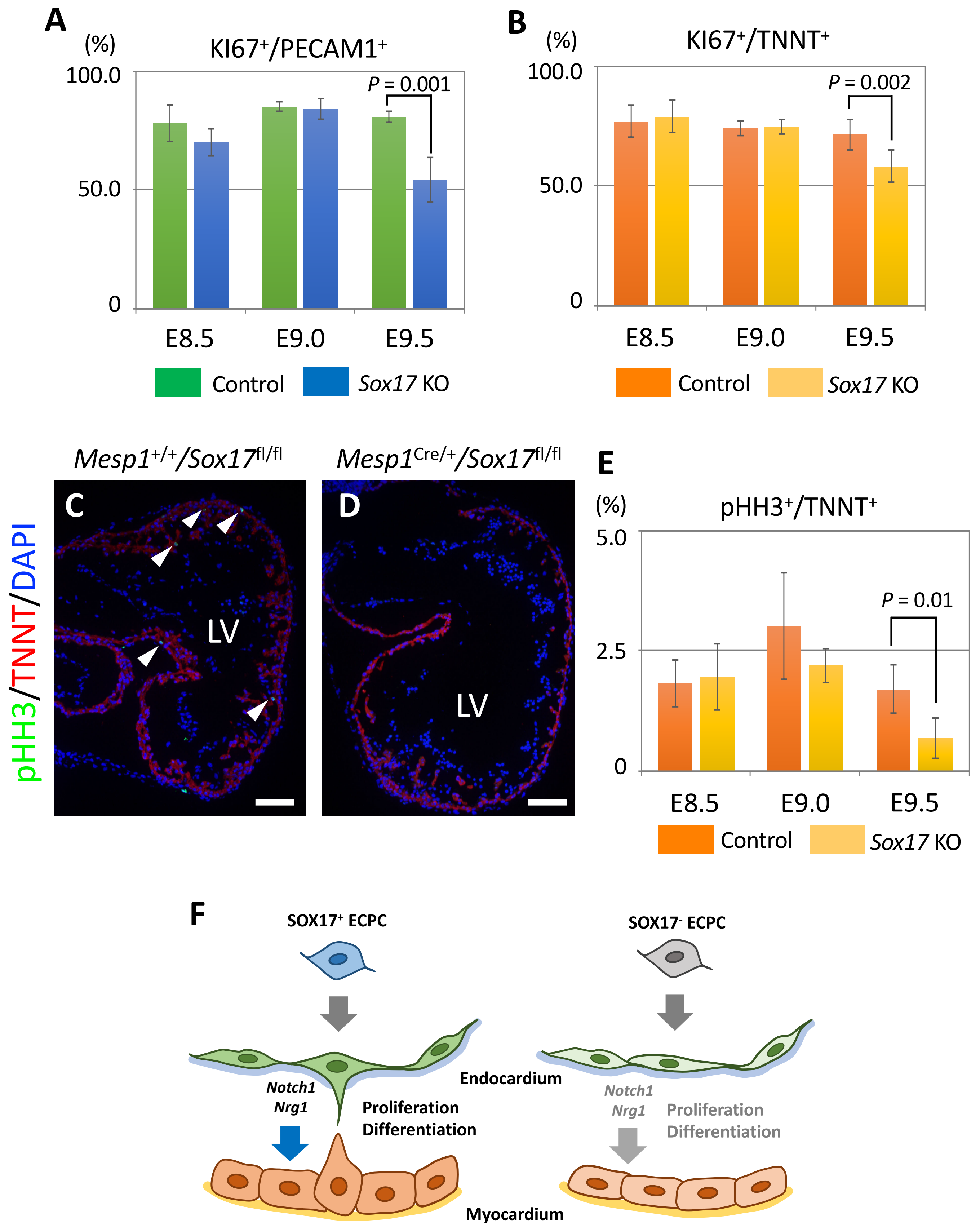
A
(%)
KI67+/PECAM1+
100.0
50.0
0
P = 0.001
E8.5
E9.0
E9.5
Control
Sox17 KO
B
(%)
KI67+/TNNT+
100.0
50.0
0
P = 0.002
E8.5
E9.0
E9.5
Control
Sox17 KO
Mesp1+/+/Sox17fl/fl
Mesp1Cre/+/Sox17fl/fl
pHH3/TNNT/DAPI
C
D
LV
LV
E
(%)
pHH3+/TNNT+
5.0
2.5
0
P = 0.01
E8.5
E9.0
E9.5
Control
Sox17 KO
F
SOX17+ ECPC
SOX17- ECPC
Endocardium
Notch1
Nrg1
Proliferation
Differentiation
Notch1
Nrg1
Proliferation
Differentiation
Myocardium
